# Intratumor Childhood Vaccine-Specific CD4^+^ T cell Recall Coordinates Antitumor CD8^+^ T cells and Eosinophils

**DOI:** 10.1101/2022.03.14.484260

**Authors:** Michael C. Brown, Zachary P. McKay, Yuanfan Yang, Georgia M. Beasley, David M. Ashley, Darell D. Bigner, Smita K. Nair, Matthias Gromeier

## Abstract

**Background:** CD4^+^ T cells are key contributors to cancer immune surveillance. However, means to effectively harness CD4^+^ T cell help for cancer immunotherapy are lacking, and antitumor mechanisms of CD4^+^ T cells remain crudely defined.

**Methods:** The impact of polio immunization on polio virotherapy was tested in syngeneic murine melanoma and breast cancer models. Antitumor effects of polio and tetanus toxoid antigens were assessed in polio and tetanus immunized mice. T and B cell knockout mice, CD4^+^ T cell adoptive transfer, and eosinophil depletion demonstrated cell-type specific contributions to the antitumor efficacy of polio recall. Phenotyping of adoptively transferred OT-I (OVA-specific) T cells in B16-OVA tumor bearing mice, as well as adoptive transfer of T cells to naïve tumor-bearing recipients, measured the impact of intratumor polio/tetanus recall on antitumor T cell immunity. Pan-cancer human transcriptome data sets were queried to test associations between eosinophils and Tregs; cytokine profiles of polio and tetanus recall were defined in human peripheral blood of healthy donors and cancer patients; CD40L blockade was used to determine dependency of recall antigen therapy on CD40:CD40L signaling.

**Results:** Prior vaccination against poliovirus substantially bolstered the antitumor efficacy of polio virotherapy in mice, and intratumor recall of poliovirus or tetanus immunity delayed tumor growth in a manner complemented by pattern recognition receptor agonist therapy and PD1 blockade. Intratumor recall antigens augmented antitumor T cell function, and caused marked tumor infiltration of type 2 innate lymphoid cells (ILC2s) and eosinophils, coinciding with decreased proportions of intratumor Tregs. Antitumor effects of recall antigens were mediated by CD4^+^ T cells, independent of CD40L signaling, and were dependent on both eosinophils and CD8^+^ T cells. Human PBMCs mounted diverse cytokine/chemokine responses, which were not impaired in patients with advanced cancer, and an inverse relationship between eosinophil and Treg signatures was observed across TCGA cancer types.

**Conclusion:** This work defines cancer immunotherapy potential of childhood vaccines, reveals their utility to engage CD4^+^ T cell help for antitumor CD8^+^ T cells, and implicates eosinophils as antitumor effectors of CD4^+^ T cells.

## INTRODUCTION

Adaptive immune memory rapidly engages anti-pathogen immunity upon repeat pathogen exposure/infection. Accordingly, recall responses, i.e. activation of adaptive memory cells upon encounter of their cognate antigen, orchestrate broad, localized innate and adaptive pro-inflammatory reactions capable of limiting replication of pathogens through antigen -dependent and -independent mechanisms^1,2^. Recent evidence implies cancer immunotherapy utility of recall responses: activating intratumor antiviral CD8^+^ T cells was shown to mediate cancer immunotherapy^3,4^.

It remains to be determined if CD4^+^ T cell recall specific to routine vaccine antigens (e.g. Poliovirus or Tetanus toxoid) hold cancer immunotherapy potential, particularly those delivered in Th2 polarizing conditions (e.g. with alum adjuvant). Despite long-standing evidence that Th1-centric immunity is optimal to mediate effective cancer immunotherapy, Th2 polarized responses hold antitumor potential^5^, eosinophil accumulation has been linked to clinical immunotherapy responses^6,7^, and type II innate lymphoid cells (ILC2s) were shown to contribute to the antitumor efficacy of PD1 blockade^8,9^. To date, much of the focus on engaging CD4^+^ T cell help for cancer immunotherapy has been on antigen-specific cancer vaccines encompassing both MHC-II and MHC-I antigens using Th1 polarized immunization strategies^10^.

PVSRIPO, the live-attenuated poliovirus type 1 (Sabin) vaccine modified with the internal ribosomal entry site (IRES) of human rhinovirus type 2^11^, has shown early evidence of efficacy in recurrent glioblastoma^12^ and recurrent, non-resectable melanoma^13,14^ after intratumor administration. Poliovirus, hereafter ‘polio’, vaccination is part of the standard pediatric immunization schedule worldwide, either with the live attenuated (Sabin), or the inactivated (Salk) vaccines. The coding sequence of PVSRIPO is identical to the type 1 Sabin vaccine. Pre-existing serum anti-polio (type 1) antibodies were confirmed in all patients receiving PVSRIPO therapy^12,13,15^. Moreover, clinical use of PVSRIPO entails prior boost with trivalent Salk (IPOL™) at least 1 week before intratumor PVSRIPO administration, which triggered increased serum anti-polio neutralizing antibody responses in all patients^12,13^. The antitumor effects of polio virotherapy encompass targeting of- and cytopathogenic damage to the neoplastic compartment and sub-lethal viral infection of monocytic lineage cells driving sustained type-I interferon signaling^15–17^. Anti-polio immunological memory, which coincides with neutralizing antibodies, is likely to impede viral replication within the tumor, but may provide an alternate antitumor mechanism of action through intratumor recall of T cells specific to polio vaccine antigens.

Using mouse tumor models of melanoma and breast cancer, we demonstrate that intratumor polio or tetanus antigens trigger local recall with marked CD4^+^ T cell, type 2 innate lymphoid cell (ILC2), and eosinophil influx; mediating antitumor efficacy through CD8^+^ T cells and eosinophils in a CD40L independent manner.

## MATERIALS AND METHODS

### Extended materials and methods are presented in Supplemental Information

#### Mice, cell lines, viruses, poly(I:C), and in vivo grade antibodies

hCD155-tg C57BL/6 mice were a gift of Satoshi Koike (Tokyo Metropolitan Institute of Medical Science, Japan). Wildtype (#000664), CD8 knockout (#002665), B cell knockout (strain code: 002288), OT-I (#003831), and CD45.1 C57BL/6 mice (#002014) were purchased from The Jackson Laboratory. OT-I mice were crossed with CD45.1 C57BL/6 mice to generate CD45.1+ OT-I mice. B16.F10 (ATCC), E0771 (G. Palmer, Duke University, USA), E0771^hCD155^, B16.F10^hCD155^, and B16.F10.9^hCD155^-OVA cells were grown in high-glucose DMEM (Gibco) containing 10% FBS. B16.F10.9^hCD155^-OVA, B16-F10^hCD155^, and E0771^hCD155^ cells were derived by lentiviral transduction with hCD155^16^. All cell lines were confirmed to be mycoplasma negative (Duke Cell Culture Facility). Laboratory grade PVSRIPO, mRIPO, and UV-inactivated PVSRIPO (UVP) were generated as previously described^15,16^. VacciGrade™ high molecular weight poly(I:C) (Invivogen) was reconstituted and annealed as recommended by the manufacturer; 30μg of poly(I:C) was delivered intratumor after mixing with either PBS or denoted antigen in the figure/figure legend. In vivo grade antibodies to IL-5 (Clone TRK5) or isotype control (Clone HRPN), PD1 (Clone RMP1.14) or isotype control (Clone 2A3), CD40L (Clone MR-1) or isotype control (catalog# BP0091), and CD40 (Clone FGK4.5) or isotype control (Clone 2A3) were purchased from BioXcell and were delivered intraperitoneal (i.p.) at the concentrations and frequencies described in figure legends.

#### Vaccines, immunizations, and intratumor viral titers

Unless otherwise indicated, polio immunization was achieved using PVSRIPO, Tetanus immunization was achieved using tetanus toxoid (Millipore-Sigma); and KLH immunization was achieved using Hemocyanin-Keyhole Limpet (Sigma-Aldrich). PVSRIPO (1×10^7^ plaque forming units/mouse), Tetanus toxoid (0.5 μg/mouse), or Hemocyanin-Keyhole Limpet (KLH, 100mg/mouse, Sigma-Aldrich) were prepared for immunization via dilution in PBS and mixed 1:1 with Alhydrogel (Invivogen). For all single antigen immunizations, 50μl of vaccine was administered bilaterally in the quadricep muscles (100μl total); combined immunizations of Salk (IPOL™) and Tenivac (Sanofi-Pasteur) were administered to only one quadricep for each vaccine (50μl), using opposing quadriceps. After the initial immunization, repeat immunization/boost occurred 14 days later. Thirty days after boost vaccination, tumors were implanted, or spleens were isolated (for adoptive transfer experiments). To determine intratumor viral titers, tumors were harvested at the denoted time points, weighed, and mechanically homogenized in 1ml PBS. Homogenate was tested by plaque assay as previously described^18^ using a starting dilution of 1:100.

#### Murine tumor model experiments

For B16 tumor implantation 2×10^5^ cells in 50μl PBS were implanted subcutaneously into the right flank; for E0771 tumor implantations 5×10^5^ cells in 50μl PBS were implanted into the 4^th^ mammary fat pad. Tumors were treated according to timelines presented in figures with either mRIPO (1×10^7^ pfu), UVP (1×10^8^ inactivated pfu), Tetanus toxoid (0.5 μg), and/or poly(I:C) (30 μg) as indicated in figures/figure legends. DMEM (dilution media for mRIPO and UVP) was injected in a similar manner for mock controls. Treatment groups were randomized by tumor volume at the first day of treatment. Tumor volume was measured by calipers and calculated using the equation (L × W × W)/2; mice were euthanized upon reaching tumor volume of 1000mm^3^. Tumor measurements were performed blinded to treatment group designations beginning after the last dose of intratumor therapy/antigen.

#### Flow cytometry analysis of tumors

Tumors were harvested at time points denoted in figures/figure legends and dissociated in RPMI-1640 media (Thermo-Fisher) containing 100μg/ml Liberase-TM (Sigma-Aldrich) and 10μg/ml DNAse I (Roche) for 30min at 37°C with agitation, followed by filtering through a 70μM (Olympus Plastics) cell strainer, centrifugation, and washing in PBS. For experiments using Zombie-Aqua, cells were stained with Zombie-Aqua in PBS (1:500) followed by washing in FACs buffer (PBS + 2% FBS) for downstream processing. Single cell suspensions were then incubated with 1:50 mouse Tru-stain FcX™ (Biolegend) in FACs buffer followed by panel-specific staining (see Supplementary Information). Flow cytometry experiments were performed using a Fortessa X20 at the Duke Cancer Institute Flow Cytometry Core Facility; FCS files were analyzed using Flow-Jo version 10 (BD Biosciences). VersaComp beads (Beckmann-Coulter) and staining of fresh blood were used to establish panel compensation, which was then optimized using florescence minus one isotype controls. Gating strategies are presented in supplemental materials; isotype controls, fluorescence minus one controls, and comparison to established negative cell populations were used to define positivity for markers of interest.

#### PBMC studies

For healthy donor analyses, leukopaks (Stemcell Technologies) from 5 different de-identified donors were processed using Leucosep™ tubes (Greiner Bio-One) and Ficoll-Paque™ Plus (GE healthcare) to isolate PBMCs following the manufacturer’s instructions. Luekopaks were collected under an approved IRB protocol held by Stemcell Technologies, Inc. PBMCs were thawed, washed in Aim-V media (Thermo-Fisher), and incubated in 1 ml AIM-V media containing 10μg/ml DNAse I (Roche). Cells were then washed in RPMI-1640 containing 10% FBS and plated at a density of 1×10^7^ PBMCs in 2ml RPMI-1640 + 10% FBS per well in a 6-well plate, followed by addition of nothing (mock), recall antigens (1μg/ml Tetanus toxoid; 1×10^8^ inactivated titer, or 50ul of IPOL™ vaccine), or Poly I:C (10μg/ml). Six days later, supernatant was retained for cytokine measurement. For PBMC studies from recurrent GBM patients, PBMCs collected under an IRB approved protocol (Duke University) and were thawed and processed as done for healthy donors, but plated at a density of 1×10^5^ cells per well in a 96 well plate and treated with mock, UVP, or Tetanus at the concentrations noted above for 3 days; a companion set of age and sex matched deidentfied PBMCs from 10 healthy donors (Stemcell Tech) were included for comparison along with batch control leukophoresis derived PBMCs. Cytokines were measured using the Human Antiviral, Human Pro-Inflammatory Chemokine, and Human Th LegendPlex™ kits (Biolegend) following the manufacturer’s instructions, and analyzed using LegendPlex software (Biolegend).

#### Adoptive transfer studies and RNA sequencing of tumor infiltrating OT-I T cells

For adoptive transfer of CD4^+^ T cells from polio or Tet immunized mice, spleens (two each) were harvested. Separately, hCD155-tg C57BL/6 mice were implanted with B16.F10.9^hCD155^ tumors six days prior to splenocyte isolation. Spleens from immunized mice were crushed through a 70μm cell strainer in 3ml RPMI-1640. Resulting cell pellets were reconstituted in 2ml ACK red blood cell lysis buffer (Lonza) and incubated for 10 min at RT, followed by addition of 10ml RPMI and centrifugation. CD4^+^ T cells were negatively selected from single cell suspensions using the EasySep™ CD4^+^ T cell isolation kit (Stemcell Technologies) following the manufacturer’s instructions. Two million CD4^+^ T cells were injected intraperitoneally six days after B16^hCD155^ tumor implantation. Seven days after tumor implantation, recipient mice were treated with mock (DMEM) or mRIPO (1×10^7^ pfu) and tumor growth was monitored. For adoptive transfer of OT-I x CD45.1 cells, spleens from OT-I x CD45.1 mice were harvested and processed to single cell suspensions as described above. OT-I splenocytes were treated with 10μg/ml SIINFEKL peptide (Invivogen) for 16h, washed in PBS, and then 2×10^6^ OT-I splenocytes were transferred in 100μl i.p. For transfer of T cells from mice treated with recall antigen therapy, splenocytes were processed as described above, washed in PBS, and transferred to naïve recipients immediately following subcutaneous injection of B16.F10.9-OVA cells. For analysis of OT-I T cell transcriptomes after polio recall, mice immunized against polio (PVSRIPO, 1×10^7^ pfu in alhydrogel) 45 days (prime) and 30 days (boost) prior to B16-OVA tumor implantation received SIINFEKL-activated OT-I splenocytes as described above, and were treated with intratumor mock (PBS) or UVP 9 days after tumor implantation. Twelve days post-treatment tumors were dissociated and subjected to CD45.1 positive selection (MojoSort mouse CD45.1 selection kit, Biolegend) per the manufacterer’s instructions. A subset of the isolated cells were used to confirm cell purity by flow cytometry for 7-AAD, CD45.2-BUV395 (BD Biosciences), CD3-PE, CD8-BV421, CD4-FITC (Biolegend unless otherwise specified); the remaining cells were lysed in 200μl Trizol (Thermo-Fisher). Cholorform extracted RNA from trizol samples (per manufacterer’s instructions) was purified using the RNEasy kit (Qiagen, Inc.) and analyzed on a Hi-seq Illumini sequencer (150bp, PE) at Azenta Life Sciences. Transcripts were aligned with STAR (v2.7) to the GRCm38 mouse genome and differential expression analysis was performed using DESeq2 (v 1.34). These isolations and downstream analyses were performed for two independent experiments: in one experiment mock treatment was compared to UVP, in a separate experiment Tet treatment was compared to UVP treatment, both in polio immunized mice.

#### Statistical analysis

Assay-specific statistical tests are indicated in the corresponding figure legends. GraphPad Prism 8 was used to perform all statistical analyses and plot data. A statistical probability of <0.05 (p<0.05) was used unless otherwise noted. All data points reflect individual specimens, experimental repeats, or mice.

## RESULTS

### Polio immunization potentiates polio virotherapy

We first tested seroconversion of mice transgenic for the human poliovirus receptor CD155 (hCD155-tg) upon immunization with trivalent Salk vaccine (IPOL) or PVSRIPO (to mimic type 1 Sabin) with and without the Th2-promoting adjuvant alum (alhydrogel; ALH), part of licensed standard vaccine formulations (e.g. Pentacel^®^, Pediarix^®^, Kinrix^®^) that enhances Salk vaccine immunogenicity^19^. A duration of 45 days post initial immunization before tumor implantation ensured sufficient time for establishment of immunological memory^20^. IPOL™ alone achieved limited seroconversion, while PVSRIPO immunization elicited a stronger antibody response; ALH bolstered antibody responses to both vaccines (**Fig S1A, B**). To recapitulate the high levels of serum neutralizing anti-polio antibodies in cancer patients prior to PVSRIPO therapy [typically >1:10,000 plaque neutralization titers^12^], we chose PVSRIPO + ALH (hereafter ‘polio’) immunization to determine the role of pre-existing polio immunity in PVSRIPO immunotherapy. Murine tumor models expressing the human poliovirus receptor CD155 and hCD155-tg mice were previously developed to permit entry and replication of PVSRIPO in neoplastic and non-malignant cells of the tumor microenvironment, and PVSRIPO was adapted to murine neoplastic host cells (mRIPO) to mediate infection and lysis^16,21^. In prior studies of mRIPO in polio vaccine naïve, immune competent mouse tumor models, a single intratumor injection failed to mediate durable antitumor responses in the absence of tumor expression of OVA or PD1/PD-L1 blockade^15,16^. However, intratumor mRIPO mediated durable antitumor effects in polio immunized mice in melanoma (B16) and breast (E0771) cancer models (**Fig 1A, B; Fig S1C**). Thus, prior polio immunization bolsters the antitumor efficacy of polio virotherapy.

**Figure 1.**
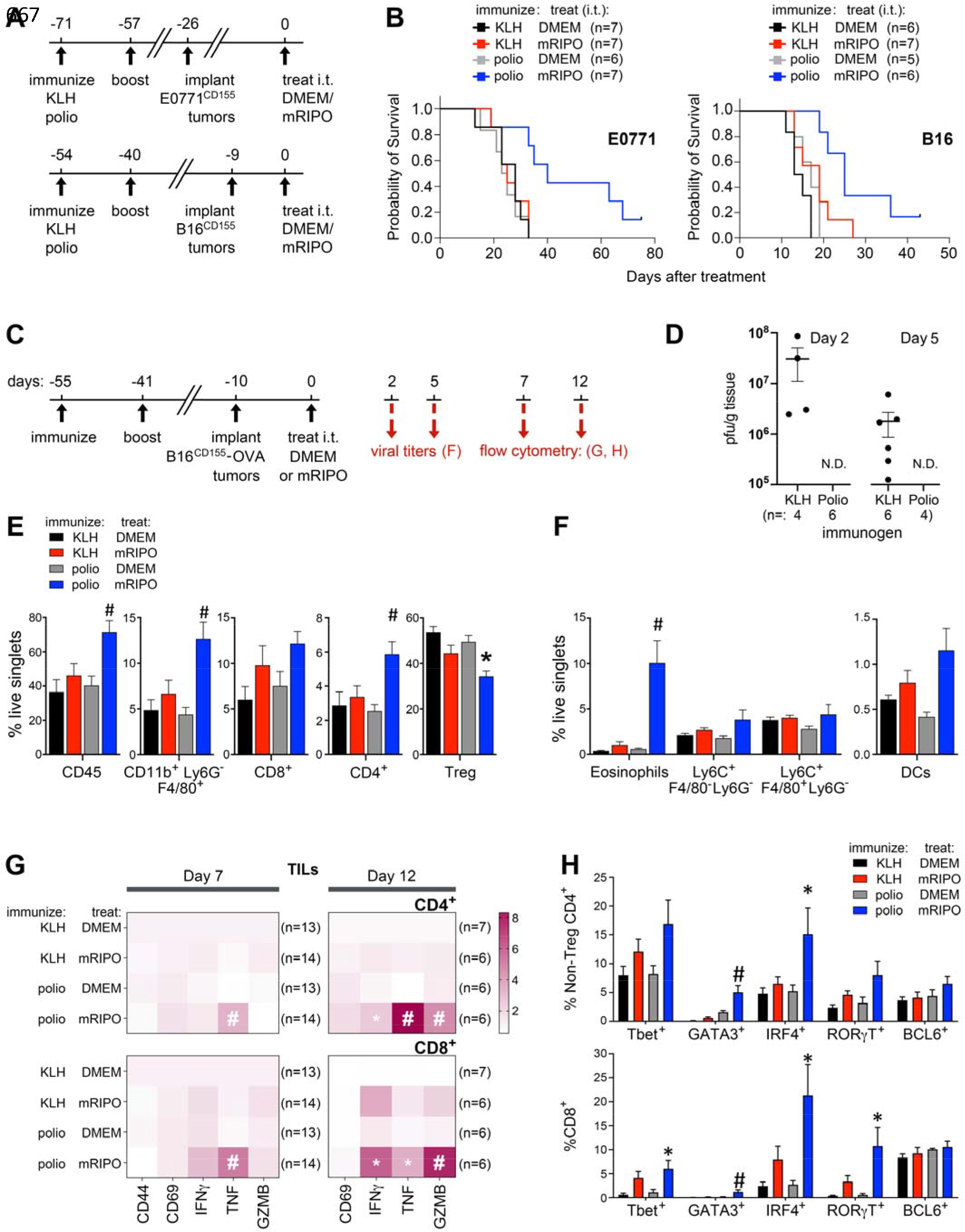
Polio immunization potentiates anti-tumor and inflammatory efficacy of recombinant poliovirus therapy. (**A**) Polio (PVSRIPO + ALH) or KLH (KLH + ALH) immunized hCD155-tg mice bearing hCD155-tg B16 (subcutaneous melanoma) or E0771 (fat pad breast cancer) tumors were treated with DMEM (control) or mRIPO (1×10^7^ pfu). (**B**) Survival was determined using the tumor volume endpoint of 1000mm^3^; see **Fig S1C** for individual tumor volumes. Representative experiments from at least two repeats are depicted. (**C**) Schema for mechanistic analyses presented in D-H. (**D**) Tumor homogenates at 2 and 5 days post-mRIPO injection were tested for the presence of infectious virus by plaque assay; N.D.= not detected. (**E**) Flow cytometry analyses of tumors 7 days after intratumor treatment for immune cell density (n=13/group mock controls; n=14/group mRIPO treated). (F) Flow cytometry analyses for myeloid cell types and dendritic cells (DCs: Ly6C^Neg^, F480^Neg^, CD11c^+^, IA/IE^+^) 11 days after treatment (n=9 KLH+Mock, n=10 KLH+mRIPO, n=5 Polio-mRIPO, n=8 polio mRIPO). (**G**) Analysis of T cell activation markers in CD4^+^ and CD8^+^ TILs at days 7 and 12 post-treatment as indicated; samples were pooled from 2 experiments. Values were normalized as fold mean KLH-DMEM values for each marker. (**H**) Intracellular staining for relevant T cell-associated transcription factors was performed on CD4^+^ (left) and CD8^+^ (right) TILs 12 days after mRIPO therapy; same samples as in day 12 of (A). Data bars indicate mean + SEM. The results were confirmed in 3 independent experiments, a representative series is shown. Figures S1-4 present extended data and gating strategies. (**E-H**) symbols indicate significant Tukey’s post-hoc test vs mock controls (*) or all other groups (#). Data bars and brackets indicate mean + SEM.

Polio virotherapy in polio vaccine naïve mice is associated with T cell inflammation within the tumor^15,16^. To determine how recall responses to polio antigens alter the tumor microenvironment (TME) after polio virotherapy we analyzed tumors at various intervals (**Fig 1C**). mRIPO replication within tumors was substantially reduced in polio immunized mice 2 and 5 days post-treatment (**Fig 1D**), consistent with high neutralizing antibody titers (**Fig S1B**). Increased total immune cell density (CD45.2^+^ cells) in polio immunized mice 7 days after mRIPO therapy was observed, explained largely by an influx of conventional CD4^+^ T cells with reduced proportions of Tregs; as well as CD11b^+^, Ly6G^Neg^, F4/80^+^ cells (**Fig 1E**). Subsequent analysis determined that the increased CD11b+ cells in polio immunized mice treated with mRIPO were eosinophils (**Fig 1F**). Neutrophil-, NK cell-, and B cell infiltration after mRIPO therapy was indistinguishable from controls (**Fig S2A**). These observations were reproducible in a separate tumor model, E0771, wherein CD11b^+^, Ly6G^neg^, F480^+^, SSC-A^hi^ cells (consistent with eosinophils) and conventional CD4^+^ T cells explained the majority of tumor-infiltrating immune cells after mRIPO therapy in polio vaccinated mice (**Fig S2B, C**).

### T cell functional phenotypes are augmented in polio immunized mice treated with mRIPO

CD8^+^ and CD4^+^ tumor-infiltrating lymphocytes (TILs) in polio vaccinated mice expressed higher levels of intracellular TNF at day 7-, and higher IFNγ, TNF, and Granzyme B at day 12 post-mRIPO treatment in the B16^hCD155^-OVA model, implying enhanced functional status (**Fig 1G**). TILs from polio immunized mice treated with mRIPO also exhibited increased differentiation phenotypes with expression of the transcription factors Tbet, GATA3, and RORγt; as well as induction of IRF4, a promoter of T cell activation and function (**Fig 1H**)^22^. Expression of the T cell exhaustion markers PD1 and TIM3 on CD4^+^ T cells in the tumor and TDLNs of polio vaccinated mice treated with mRIPO were reduced (**Fig S2D**). Changes in T cell activation/ differentiation markers were consistent in the E0771 orthotopic breast cancer model (**Fig S2E, F**). Collectively, these findings imply that recall responses to polio potentiate T cell function after intratumoral polio virotherapy.

### Polio and tetanus recall antigens mediate antitumor efficacy and induce broad inflammatory responses in human PBMCs

Recent work indicates that reovirus-specific memory CD8^+^ T cells directly kill cancer cells to potentiate oncolytic reovirus therapy^23^, and that tetanus-specific memory CD4^+^ T cells kill pancreatic cancer cells infected with *Listeria* vector expressing tetanus toxoid^24^. Alternatively, inflammatory responses caused by intratumor memory T cell recall in the tumor microenvironment may also bolster immune surveillance, as shown for antiviral memory CD8^+^ T cell antigens (e.g. CMV)^3,4^. We hypothesized that recall-induced inflammation explained accentuated antitumor efficacy of polio virotherapy in polio vaccinated mice, since viral replication—required for production of viral antigen in tumor cells, oncolysis, and antiviral innate inflammation^15^—was dramatically reduced in polio immunized mice (**Fig 1D**). To probe the antitumor effects of polio recall in the tumor microenvironment we used a model devoid of hCD155 (mice and tumors) and treated tumors with UV-inactivated PVSRIPO (‘UVP’) to preclude tumor cell/TME infection and viral replication. We included comparisons with another vaccine-associated recall antigen, Tetanus toxoid (Tet).

Intratumor therapy with UVP exerted antitumor efficacy exclusively in polio immunized mice; Tet treatment mediated a transient antitumor effect only in Tet immunized mice (**Fig 2A, B; Fig S5C**). We confirmed these effects in mice co-immunized with clinical grade Tenivac (tetanus and diphtheria) and IPOL: intratumor therapy with UVP and Tet mediated antitumor efficacy exclusively in Tenivac + IPOL immunized mice (**Fig 2C**). Natural recall responses typically occur in the presence of a localized innate immune response to pathogen replication. To mimic this, Tet or polio immunized mice were treated with high molecular weight poly(I:C) alone or in combination with UVP or Tet. Both UVP and Tet mediated pronounced antitumor effects in this context (**Fig 2D**). To gauge the inflammatory potential of Polio and Tetanus antigens in a human system, we measured cytokine responses to UVP, Tet, and IPOL relative to poly(I:C) (positive control) after 6 days (**Fig 2E**) in peripheral blood mononuclear cells (PBMCs), wherein CXCR3 ligands (CXCL9, 10, 11), IFNs (α/β/λ/γ), and other cytokines were induced. PBMCs from patients with recurrent GBM mounted similar responses to UVP and Tet (**Fig 2E, Fig S5D-F**), indicating that immunological memory to tetanus and polio vaccines are not compromised in patients with heavily pre-treated, advanced cancer. Together, these observations indicate potential of recall antigens in inducing antitumor effects.

**Figure 2.**
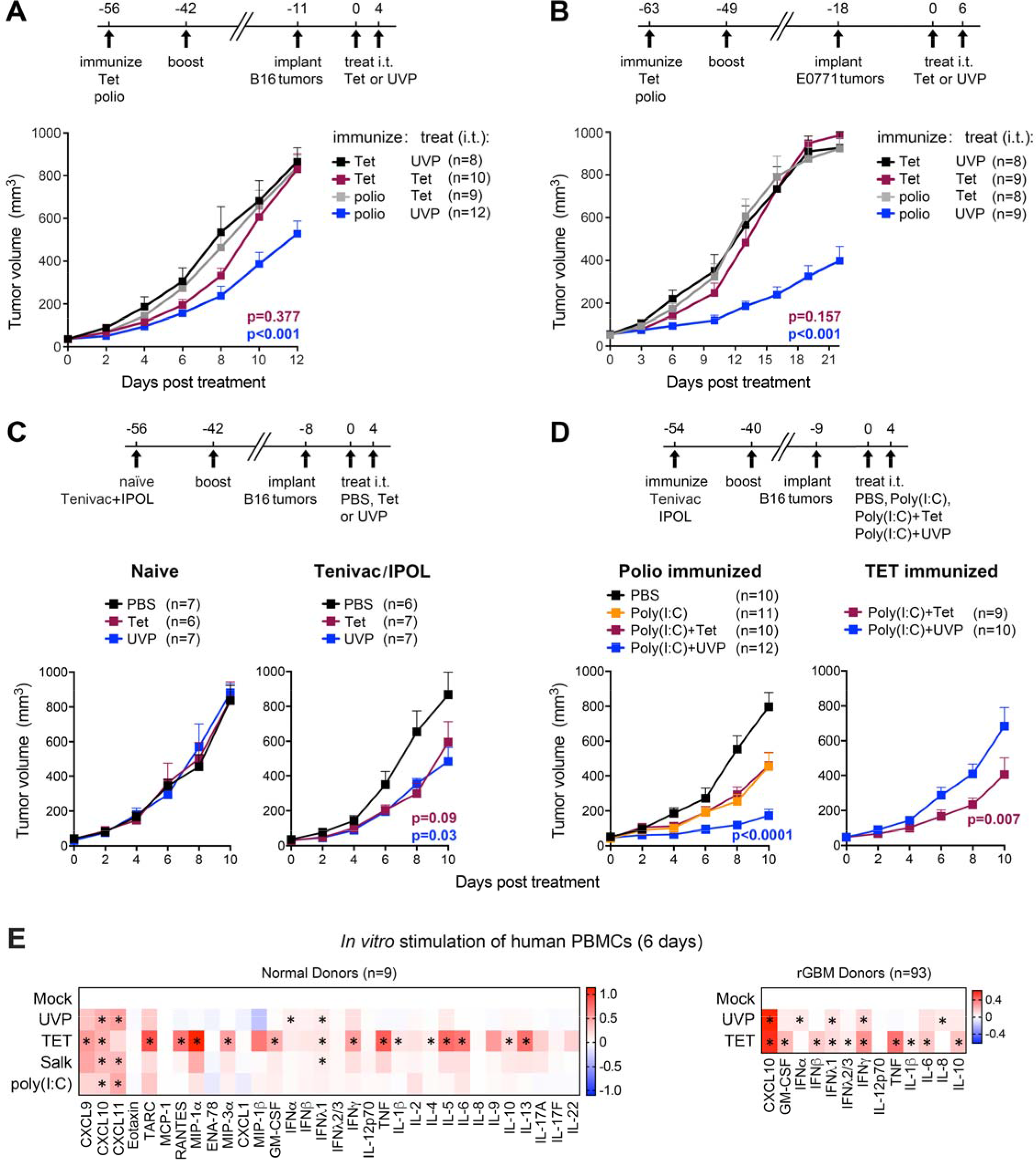
Polio and tetanus recall antigens mediate antitumor efficacy and induce broad inflammatory responses in human PBMCs. B16 (**A**) or E0771 (**B**) tumor-bearing mice immunized with polio (PVSRIPO + ALH) or Tet (Tetanus toxoid + ALH) were treated intratumor with Tet (tetanus toxoid) or UVP (UV-inactivated PVSRIPO) as shown. (**C**) Age-matched naïve or Tenivac and IPOL™ immunized mice were treated with PBS, Tet, or UVP as shown. (**D**) Mice immunized as in (A, B) were treated intratumor with mock, poly(I:C) (30μg), poly(I:C) + Tet, or poly(I:C) + UVP as shown. (**E**) Human PBMCs from healthy donors (n= 9 tests from 5 different donors) or recurrent GBM patients were treated with mock, UVP (1×10^8^ inactivated), or Tet (1μg/ml) for 6 or 3 days, respectively, *in vitro*. Supernatant cytokines were measured and are presented as log-fold mock control; asterisks denote significant Tukey’s post hoc test. (**A-D**) Mean + SEM are shown and a representative experiment of at least two repeats is shown; asterisks indicate Dunnett’s multiple comparison test p<0.05 (two tailed) vs all other groups. **Figure S5** presents related data.

### CD4^+^ T cells mediate the antitumor efficacy of recall antigens

To determine which adaptive compartment(s) mediate(s) antitumor efficacy of recall responses, we compared intratumor treatment with UVP in CD8^+^ T and B cell knockout (k/o) mice relative to wildtype (wt) mice (**Fig 3A**). We did not address this question in CD4 k/o mice due to the anticipated role of CD4^+^ T cells in influencing both CD8^+^ T cell and B cell responses to vaccination, rendering the comparison uninformative. Polio immunization in wt and CD8 k/o mice led to anti-polio antibody production; as expected, sera from polio immunized B cell k/o mice did not react to polio capsid (**Fig 3B**). B16 tumor growth was similar in each genetic context after mock treatment. Relative to wt mice, the antitumor efficacy of UVP in polio immunized mice was limited in CD8 k/o mice at later time points, and was enhanced in B cell k/o mice (**Fig 3C**). Eosinophils express Fcγ receptors, contribute to antibody dependent cellular cytotoxicity, and may be influenced by antibody/ complement-mediated activites^25^. To test if eosinophil influx or other changes observed in the TME after polio recall (**Fig 1**) are dependent on B cells (or antibodies), we analyzed tumor infiltrating cells seven days after treatment in polio immunized wt vs B cells k/o mice after mock or UVP treatment. Similar to polio virotherapy in polio immunized mice, UVP treatment led to increased immune cell influx, eosinophil infiltration, along with increased CD4^+^ T cells—decreased proportions of which were Tregs (**Fig 3D**). Thus, the antitumor efficacy of polio recall is independent of, and possibly countered by, B cells, and partially dependent upon CD8^+^ T cells.

**Figure 3.**
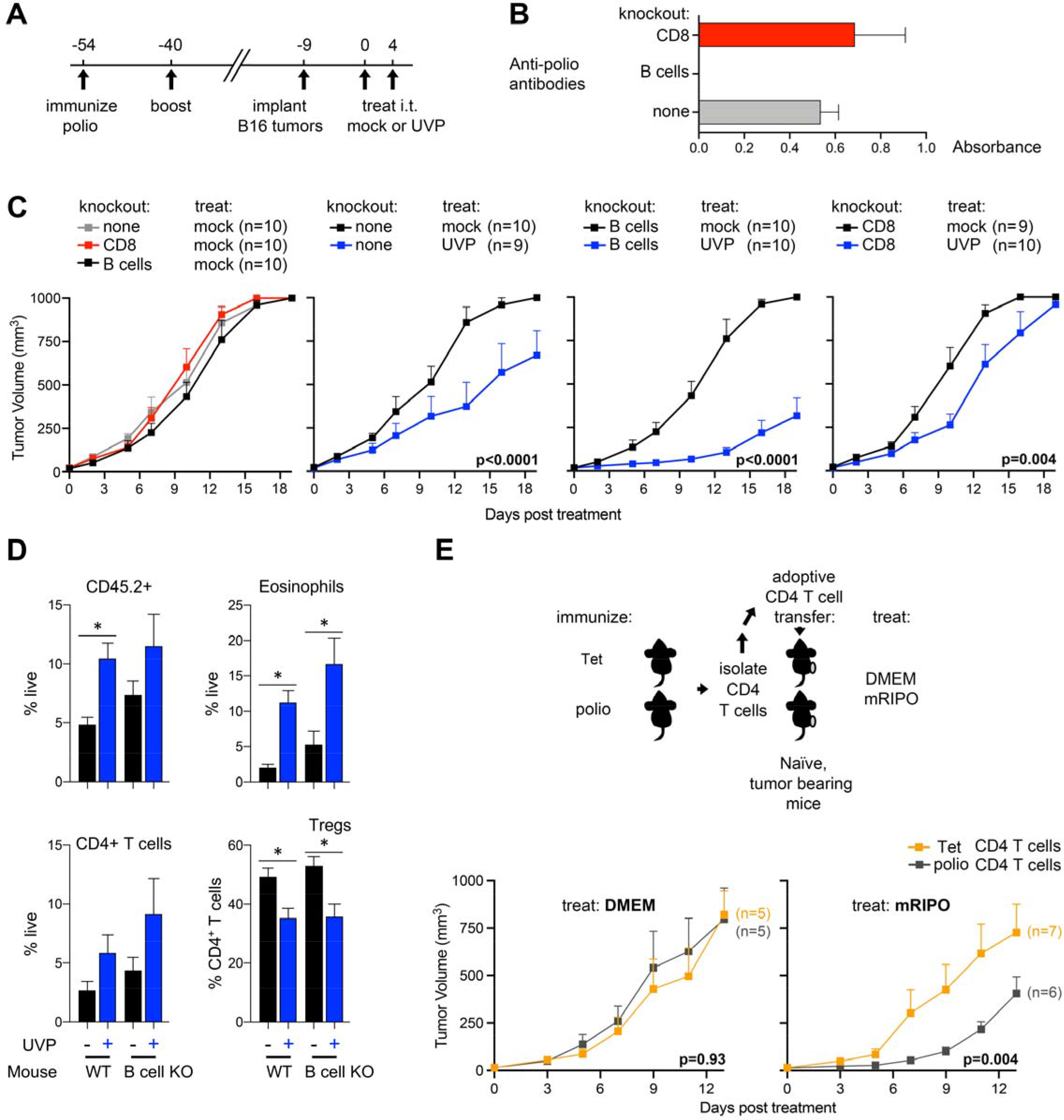
CD4^+^ T cells mediate antitumor efficacy of polio recall. (**A**) Schema depicting experimental design for tumor growth experiments in wt, CD8^+^ T cell k/o, or B cell k/o mice presented in (B, C). (**B**) ELISA for serum anti-polio antibodies in animals of each genetic background at the time of tumor implantation (n= 4/group). (**C**) Mean tumor volume + SEM after mock treatment (left) or mock vs UVP treatment (right panels) for each genotype context; p-values are from two-way ANOVA comparison. (**D**) Flow cytometry analysis of tumors seven days after mock or UVP from wt or B cell k/o mice immunized and treated as in A; mean+SEM is shown and asterisks denote two-tailed tukey’s post hoc test p<0.05. (**E**, top) CD4^+^ T cells from spleens of mice immunized with Tet (control) or polio as in Figure 2A were adoptively transferred to naïve B16 tumor-bearing recipients 1 day prior to intratumoral treatment with DMEM or mRIPO. (**E**, bottom) Mean tumor volume + SEM for mock or mRIPO treated mice for each CD4^+^ T cell transfer condition; p-values are from two-way ANOVA comparison to the control group. See Figure S6 for a related repeat data set.

These observations, along with robust CD4^+^ T cell infiltration in tumors after mRIPO therapy (**Fig 1E**), suggest that CD4^+^ T cells dictate the antitumor efficacy of polio recall responses. To test if polio-specific CD4^+^ T cell recall responses are sufficient to potentiate the antitumor efficacy of mRIPO therapy, we isolated CD4^+^ T cells from polio or Tet immunized mice, transferred them to naïve recipients bearing B16 tumors, and treated tumors with DMEM (control) or mRIPO. The antitumor efficacy of mRIPO was markedly enhanced in mice with CD4^+^ T cell adoptive transfer from polio immunized mice relative to that of Tet immunized mice (**Fig 3E**). We repeated this experiment comparing transfer of CD4^+^ T cells from spleens of KLH vs. polio immunized mice (**Fig S6**). Similarly, the antitumor efficacy of mRIPO was enhanced after transfer of CD4^+^ T cells from polio vaccinated mice. Thus, CD4^+^ T cells mediate the antitumor efficacy of polio recall.

### Intratumor recall antigen therapy potentiates antitumor CD8^+^ T cell function

Tumor-specific CD4^+^ T cells were shown to mediate antitumor efficacy through direct tumor cell killing^26–28^, engagement of cytotoxic innate immune cells^29,30^, or by providing CD4^+^ T cell help to effector CD8^+^ T cells^31–34^. Our data demonstrate that recall antigens not expressed by tumor cells (i.e. vaccine associated antigens), also engage antitumor functions of CD4^+^ T cells when delivered intratumorally (**Figs 2–3**). The lack of expression and thus, MHC class II presentation, of recall antigens in cancer cells within this system precludes the contribution of direct antigen-specific cancer cell killing by cytolytic CD4^+^ T cells. Rather, mRIPO therapy in polio vaccinated mice was associated with enhanced polyfunctional CD8^+^ T cell phenotypes (**Fig 1G**), and the antitumor effect of UVP in such mice was tempered in mice lacking CD8^+^ T cells (**Fig 3C**). Thus, we next sought to determine if polio (UVP) and Tet recall enhances the function of antitumor CD8^+^ T cells. To this end we adoptively transferred CD45.1^+^ marked, *ex vivo* SIINFEKL-stimulated OT-I CD8^+^ T cells to polio or Tet vaccinated mice and determined the impact of UVP and Tet-induced recall on B16-OVA tumor infiltrating OT-I T cell phenotype (**Fig 4A**). As in prior studies, induction of recall responses in the tumor after UVP (in polio immunized mice) or Tet (in Tenivac immunized mice) was associated with delayed tumor growth and increased tumor infiltration of CD45^+^ cells, eosinophils, and conventional CD4^+^ T cells (**Fig 4B**). A non-significant increase in OT-I and endogenous CD8^+^ TILs was also observed (**Fig 4B**). Analysis of tumor infiltrating OT-I CD8^+^ T cells revealed enhanced Granzyme B, TNF, and IFNγ after UVP treatment in polio immunized mice or Tet treatment in Tenivac immunized mice (**Fig 4C**). As observed in polio vaccinated mice after mRIPO therapy (**Fig 1G, H**), transcription factors associated with varied Th polarizations were induced in both OT-I and endogenous T cells, including GATA3, RORγT, and BCL6 (**Fig 4C**). To assess changes in antitumor T cells after intratumor polio recall in an unbiased manner, tumor infiltrating OT-I T cells were isolated after mock or UVP treatment of polio immunized mice and their transcriptomes were analyzed by RNA-sequencing (**Fig 4D, Fig S8**). Downregulated expression of genes associated with NF-κB and SRC signaling (*S100b, RSAD2, Mpzl1*) coincided with increased expression of granzymes; genes linked with T cell activation, function, or homeostatis (*Taok3, CD86, CCR5, Egr2, Adgre1, Vdr, IRF4*, and *BCL6*); genes associated with Th1 immunity (*Ptger4, Fgl2*); as well as genes associated with Th2 immunity (*Alox15, Ccl8*, and *GATA3*) (**Fig 4D**, **Fig S8C**). Demonstrating enhanced immunological memory against B16-OVA tumors after intratumor recall, T cells isolated from spleens of mice in which intratumor recall occurred (UVP in polio vaccinated, or Tet in Tenivac vaccinated) delayed tumor growth upon transfer to mice implanted with B16-OVA tumors (**Fig 4E**). Collectively, these observations indicate that intratumor recall antigen therapy potentiates the function of antitumor T cells with both Th1- and Th2-associated attributes.

**Figure 4.**
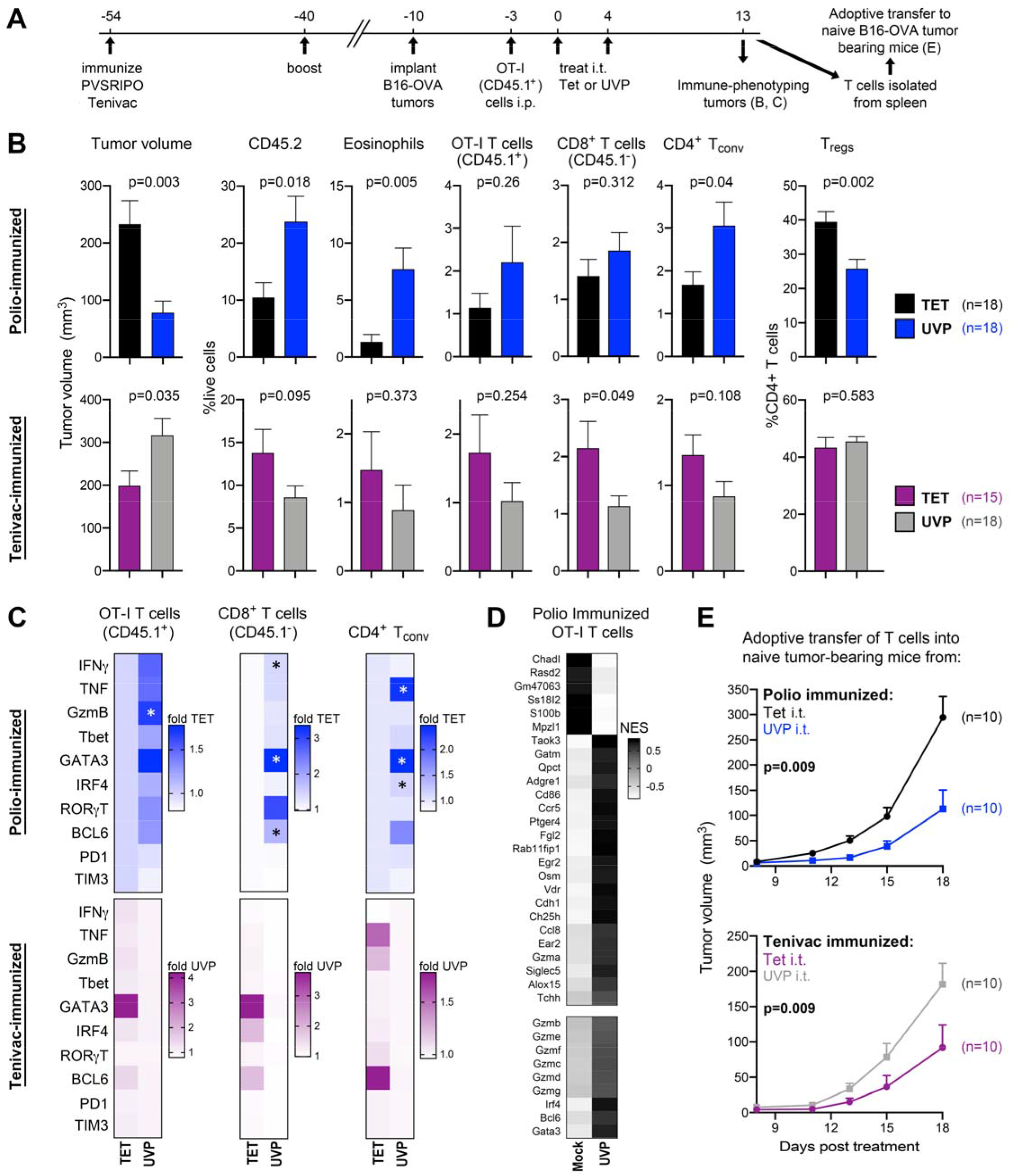
Intratumor recall antigen therapy potentiates antitumor CD8^+^ T cell function. (**A**) Mice immunized against polio (PVSRIPO) or Tet (Tenivac) were implanted with B16-OVA tumors, followed by adoptive transfer of ex vivo activated OT-I (CD45.1^+^) cells, and treatment with either Tet or UVP. Thirteen days after treatment tumors and spleens were harvested for analyses in (**B-D**). (**B**) Tumor volume and flow cytometry analysis from mice treated as in (A). (**C**) Analysis of TIL subsets (OT-I, endogenous CD8^+^, and conventional CD4^+^ T cells) for markers of activation and differentiation; gating for OT-I cells is presented in Figure S7. (**D**) Tumor infiltrating OT-I T cells were isolated from polio immunized mice treated with either mock or UVP as in Fig 4A 12 days after treatment. RNA was isolated from CD45.1+ enriched OT-I T cells (see Fig S8A for validation) and analyzed by RNA sequencing. Center and scaled mean enrichment scores are shown for genes that were significantly enriched or depleted in two independent experiments (top panel) or for granzymes, IRF4, BCL6, and GATA3 (bottom panel); n=4 replicates/group. Fig S8B-C presents False Discovery Rate corrected p-values, sample-level enrichment scores from 2 independent experiments, and annotation of relevance of significantly enriched/depleted transcripts. (**E**) Tumor progression in naïve mice upon adoptive transfer of T cells from spleens of mice from experiment in A-C. Mean tumor volume + SEM is shown; p-value is from two-way ANOVA. All data bars represent mean + SEM; heatmaps in (C) were normalized by fold average of the mismatched antigen control; symbols indicate significant Tukey’s post-hoc test relative to all other groups (#) or respective DMEM control (*). (A-C) represents pooled results from two independent experiments, data in (E) were repeated twice and a representative series is shown.

### Tumor infiltrating eosinophils inversely associate with Tregs in human tumors

In addition to increased T cell functional phenotypes and CD4^+^ T cell infiltration coinciding with reduced proportions of Tregs, polio and tetanus recall was associated with a marked increase in tumor infiltrating eosinophils (**Figs 1, 3 and 4**). Tumor eosinophil influx has been shown to be associated with immunotherapy response^6,7,9^, eosinophils can contribute to immune surveillance^35^, and vaccination with GMCSF expressing tumor cells contributed to antitumor effects of CD4^+^ T cell help in an IL-5 dependent manner^36^. To query whether a natural relationship exists between eosinophils and the tumor immune microenvironment composition in human tumors, we analyzed a pan-cancer TCGA data set^37^ for relationships between eosinophils and other immune cell types. Using previously computed CIBERSORT^38^ prediction of cell infiltrates^37^, samples from each cancer type were stratified by presence or absence of detected eosinophil gene expression signatures (**Fig 5A, Figs S9-10**); TGCT and UVM were excluded from downstream analyses due to limited numbers of cases with eosinophil enrichment (n < 3). Low grade gliomas (LGG) had the highest eosinophil enrichment across all tested tumor types, with most tumor types having between 5 - 20% of cases with eosinophil signatures (**Fig S9A**). While no significant association of eosinophil presence was observed with CD8^+^ or CD4^+^ T cell enrichment, eosinophil presence was associated with significantly lower Treg, but increased monocyte signatures across all cancer types (**Fig 5A, Fig S9B-C**). The detection of eosinophils was associated with longer survival in LGG, but not other tumor types (**Fig 5B, Fig S9D**). Significant differences in Treg density upon stratification by eosinophil enrichment were observed within several cancer types, with heterogenous relationships between CD4^+^ and CD8^+^ T cell density (**Fig S10A**). In melanoma (SKCM) and colon adenocarcinoma (COAD), eosinophil presence was associated with significant CD4^+^ T cell enrichment (**Fig S10B**). These data may reflect a role for eosinophils in influencing tumor infiltrating T cell biology, in particular, Tregs.

**Figure 5.**
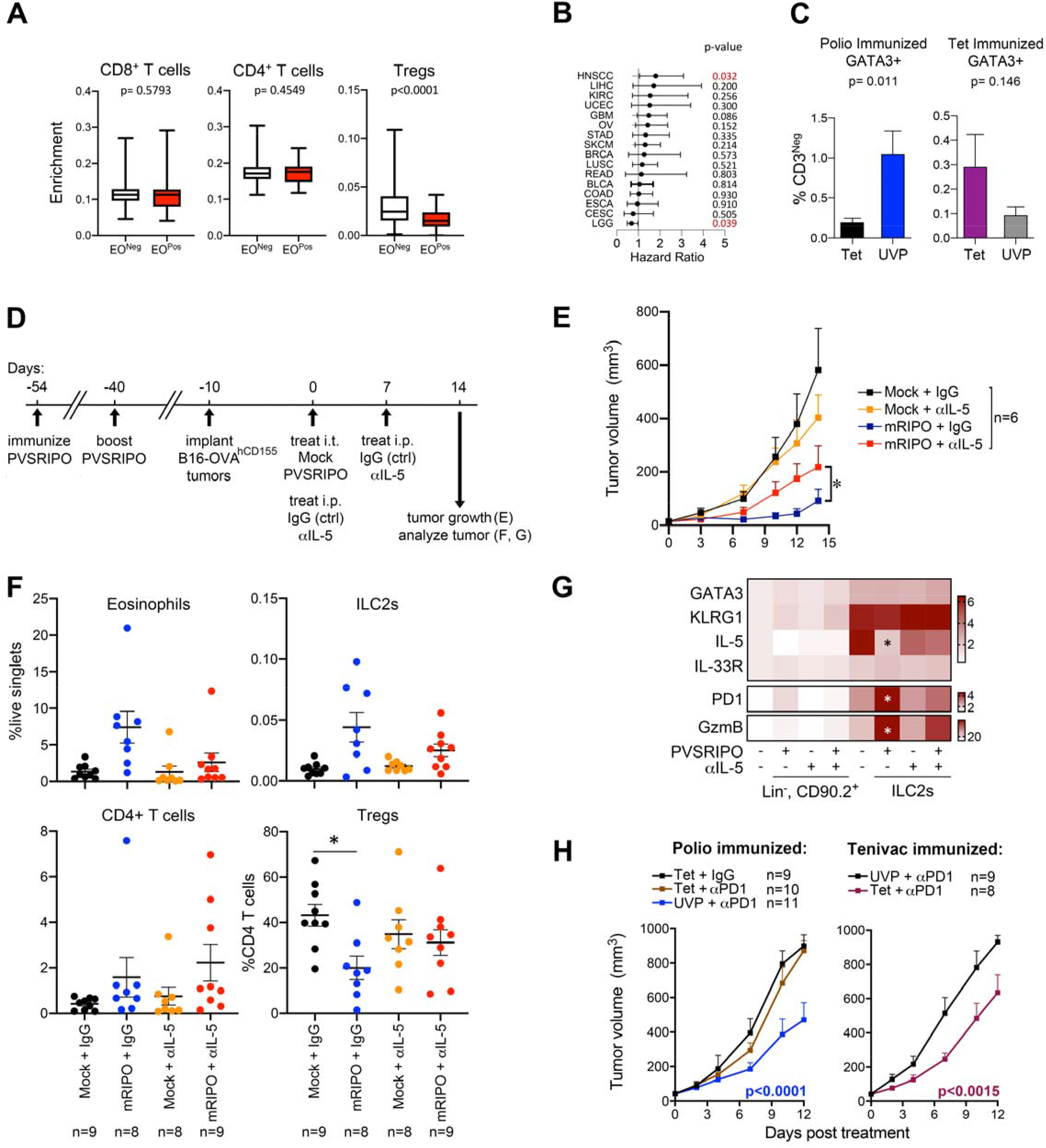
Eosinophils inversely associate with Tregs in human tumors; mRIPO induces antitumor antitumor type II immunity in polio immunized mice. (**A**) Enrichment scores of CD8^+^ T cells, CD4^+^ T cells, or Tregs in 29 different cancer types, p-values are from paired t-test. (**B**) Hazard ratios −/+ 95% confidence intervals for denoted cancer types; cancer types with less than 20 deaths in the cohort were excluded for these analyses; p-value is from Mantel-Cox log-rank test. (**C**) Percentage of GATA3+ CD3^Neg^ cells re-gated from flow cytometry data set presented in Fig 4C, p-values are from unpaired t-test. (**D**) Design for (E-H): hCD155-transgenic mice were vaccinated against polio, implanted with B16-OVA^hCD155^ tumors, and treated with intratumor mock or PVSRIPO in the presence or absence of eosinophil depletion (anti-IL-5 antibody or IgG control, 1mg administered once weekly). Mean tumor volume + SEM (**E**), eosinophil and ILC2 density in tumors (**F**), and phenotype of tumor infiltrating ILC2s (lineage^Neg^, CD90^+^, CD127^+^, CD25^+^) vs lineage negative CD90^+^ cells for comparison are shown (**G**). (**H**) Polio or tetanus immunized mice were treated i.t. with Tet or UVP with control IgG (250μg/mouse every 3 days) or in combination with anti-PD1 antibody (250μg/mouse every 3 days); mean tumor volume + SEM is shown. (**E**) Asterisk indicates Two-way ANOVA p<0.05 (two-tailed); (**F-G**) asterisk indicates Tukey’s post hoc test p<0.05 relative to mock + IgG control; (**H**) p-values are from two-way ANOVA p<0.05 compared to recall antigen control. Figs S9-S11 presents gating strategy and extended data.

### Antitumor type II immunity after mRIPO treatment of polio immunized mice

Eosinophil recruitment can be mediated by innate lymphoid type 2 cells (ILC2s), which coordinate Th2 responses through direct interactions with CD4^+^ T cells^39^, express the transcription factor GATA3 and eosinophil recruiting cytokine IL-5, but lack T cell receptor expression. In gating from prior experiments originally designed to detect GATA3 expression in T cells (**Fig 4B-C**), we noted increased GATA3^+^ CD3^Neg^ cells in tumors of mice after triggering intratumor recall with either polio or tetanus antigen (**Fig 5C**), possibly reflecting ILC2 influx. We next tested how eosinophils impact PVSRIPO therapy in polio immunized mice, and measured changes in ILC2s directly. Polio immunized, B16-OVA^hCD155^ tumor bearing mice with and without concomitant eosinophil depletion [via anti-IL-5 antibody^40^] were treated with mock or mRIPO (**Fig 5D, E**). Eosinophil depletion, which was confirmed in blood and tumors 14 days after treatment (**Fig 5F, Fig S11C**), mitigated the antitumor effects of polio recall (**Fig 5E**). Depletion of eosinophils did not reduce CD4^+^ T cell influx after PVSRIPO treatment in polio immunized mice, but blocked reductions in Treg proportion (**Fig 5F**). Thus, eosinophils are associated with decreased Treg density in human tumors (**Fig 5A**), and suppress Treg populations during recall antigen therapy. Moreover, intratumor PVSRIPO therapy led to ILC2 influx (**Fig 5F**) and altered ILC2 phenotypes, with reduced IL-5 and induced PD1 and Granzyme B expression (**Fig 5G**). These data demonstrate antitumor roles for eosinophils and recruitment of ILC2s with altered phenotypes after recall antigen therapy. Recent work indicated that PD1-expressing ILC2s contribute to immune checkpoint blockade therapy^8,9^. Possibly due in part to PD1-expressing ILC2s, recall antigens potentiated anti-PD1 therapy in mice (**Fig 5H**).

### The antitumor efficacy of polio recall is independent of CD40L-CD40 signaling

A key mechanism by which CD4^+^ T cell help mediates antitumor efficacy is through CD40L signaling to CD40 on antigen presenting cells^31^. Indeed, agonistic antibodies to CD40 are being tested as cancer immunotherapy agents^41^. To determine if recall antigens mediate antitumor effects through CD40 signaling, we compared the antitumor efficacy of polio recall with and without CD40L blockade (**Fig 6A, B**) in the context of OT-I CD8^+^ T cell transfer as in Figure 5. While a trend towards more aggressive tumor growth was observed after CD40L blockade, it did not affect antitumor effects of UVP or UVP-elicited influx of ILC2s, eosinophils, conventional CD4^+^ T cells, or antitumor OT-I T cells (**Fig 6B**). Moreover, an agonistic anti-CD40 antibody did not recapitulate ILC2 and eosinophil influx observed after recall antigen therapy, but was instead associated with an increase in tumor associated macrophage density (**Fig 6C, D**). Thus, we conclude that intratumor CD4^+^ T cell recall engages antitumor type I and II immunity independent of CD40-CD40L signaling.

**Figure 6.**
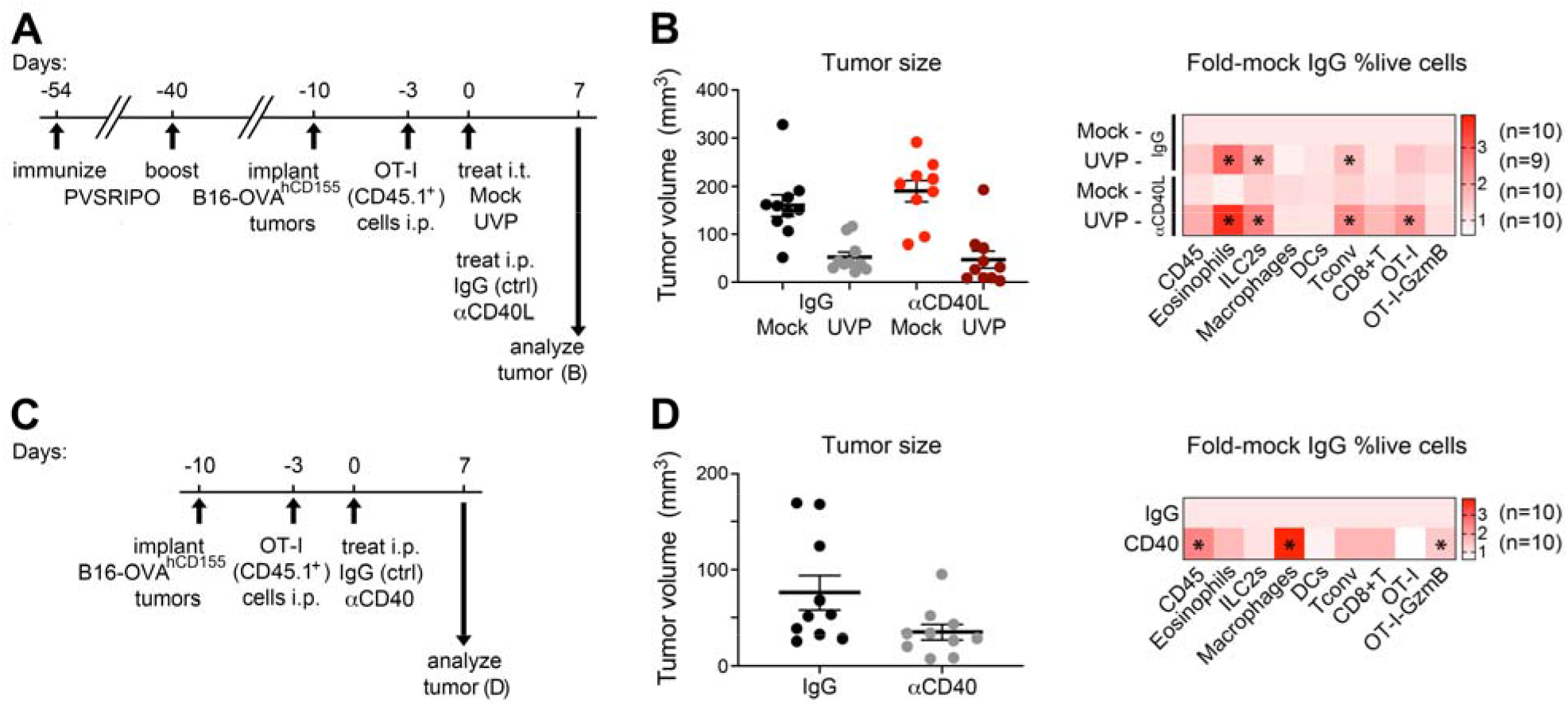
The antitumor efficacy of recall antigens is independent of CD40L-CD40 signaling. (**A**) B16-OVA tumor bearing mice previously immunized against polio received adoptive transfer of OT-I T cells followed by intratumor treatment with either mock or UVP concomitant with control IgG or a CD40L blocking antibody (250μg, every 3 days) for experiment shown in (B). (**B**) Tumor volume (left) and analysis of tumor infiltrating immune cells (right) was conducted 7 days after treatment. Data bars depict mean −/+ SEM; heatmap depicts fold-mock IgG control. (**C**) B16-OVA tumor bearing mice received adoptive transfer of OT-I cells followed by intraperitoneal treatment with control IgG or anti-CD40 antibody (50μg/mouse, once) for experiment shown in (D). (**D**) Tumor volume (left) and analysis of tumor infiltrating immune cells (right) were measured 7 days after treatment. Data bars depict mean −/+ SEM; heatmap depicts fold-mock IgG control. Asterisks indicate Dunnett’s post-hoc test relative to mock + IgG control (**B**), or unpaired t test relative to IgG control (**D**). All experiments were repeated at least twice and representative series are shown.

## DISCUSSION

This work reveals cancer immunotherapy potential of tumor-localized CD4^+^ T cell recall, engaging multifaceted inflammatory cascades involving both innate and T cell compartments. CD8^+^ T cells and eosinophils contributed to the antitumor efficacy of polio recall/virotherapy in vaccinated mice, indicating that CD4^+^ T cell recall coordinates multiple antitumor effectors.

CD4^+^ T cell help is key for generating fully functional antitumor CD8^+^ T cell immunity^10,31^ and long-term memory CD8^+^ T cells^33,42^. Antitumor CD8^+^ T cells exhibited greater polyfunctional phenotypes after recall response induction, adoptive T cell transfer from recall antigen treated mice delayed tumor growth in naïve recipients, and UVP antitumor effects were blunted in polio immunized mice lacking CD8^+^ T cells. This indicates provision of CD4^+^ T cell help to antitumor CD8^+^ T cells during polio virotherapy and/or injection of recall antigen into tumors. Indeed, CD4^+^ T cell help is linked to T cell exhaustion marker downregulation, increased TNF/IFNγ/Granzyme B expression in effector T cells, and Tbet/IRF4 induction in ‘helped’ effector T cells^32^, all of which were observed after polio virotherapy in polio immunized mice.

Such T cell help likely involves multiple mechanisms, including enhanced activation of tumor antigen presenting dendritic cells; CD4^+^ T cell secretion of CD8^+^ T cell supportive cytokines like IL-21 and IFNγ; TME reprogramming to a CD8^+^ T cell supportive environment, e.g. via IFNγ secretion to induce MHC-class I antigen presentation; or through positive effects of eosinophils on CD8^+^ T cell immune surveillance^35^. Antitumor efficacy of polio recall in the context of CD40L blockade and the failure of CD40 agonism to recapitulate recall antigen therapy phenotypes show that the antitumor effects of recall antigens do not rely entirely on CD40L signaling.

The antitumor efficacy of polio recall only partially depended on CD8^+^ T cells. Recall antigen therapy generated robust eosinophil and ILC2 influx, and eosinophil depletion decreased antitumor effects in polio immunized mice treated with mRIPO. In asthma, CD4^+^ T cells have been shown to recruit eosinophils^43^ via ILC2s^44^. ILC2s express MHC-class II and interact with CD4^+^ T cells to propagate Th2 responses in helminth infections^39^. In cancer, eosinophil infiltration is linked with immunotherapy response^6,7^, eosinophils were shown to support CD8^+^ T cell immune surveillance^35^, and ILC2s express PD1 and contribute to anti-PD1 antitumor efficacy^8,9^.

We discovered an inverse relationship between tumor eosinophil influx and Treg density in human cancers, possibly reflecting a role for eosinophils in Treg suppression. Indeed, recall antigen therapy led to decreased proportions of Tregs that covaried with eosinophil influx/IL-5. Intringuingly, Treg depletion leads to increased eosinophil influx in murine tumor models^45^ and decreased Treg proportions and function is observed in asthmatic lung tissue^46^ which, together with our observations imply reciprocal antagonism between Tregs and eosinophils. We also observed increased PD1 and Granzyme B expression in tumor-infiltrating ILC2s after polio recall, possibly indicating their transition to an antitumor phenotype amenable to PD1 blockade. Recall antigen therapy accentuated the antitumor efficacy of PD1 blockade despite stagnant or reduced PD1 expression on T cells. Varying ILC2 phenotypic states have been observed, including transition to “Th1-like” polarity^47^. Highlighting the pivotal importance of context, both eosinophils and ILC2s have also been shown to mediate pro-tumorigenic effects^48,49^.

We used Th2 polarizing vaccination strategies, consistent with the clinical use of polio and tetanus vaccines, to decipher antitumor potential of CD4^+^ T cell recall. Based on studies revealing the utility of CD8^+^ T cell recall in mediating antiviral inflammation^1,2^ and cancer immunotherapy^3^, we hypothesize that recall of Th1-polarizing vaccines will also parlay help to antitumor CD8^+^ T cells, with the caveat that antitumor functions of type II immunity may be lacking. While—canonically—Th1/Tc1, Th2/Tc2, and Th17/Tc17 polarizations are mutually exclusive, recall antigen therapy produced Th1 (CD8^+^ T cell engagement, expression of Tbet/IFNγ by CD4^+^ T cells); Th2 (eosinophil/ILC2 recruitment, expression of GATA3 by CD4^+^ T cells); and to a lesser extent, Th17 (RORγT expression in CD4^+^/CD8^+^ T cells) polarizing features. Thus, diverse CD4^+^ T cell polarizations can generate compatible and productive immune responses^50^.

Our work demonstrates that while pre-existing immunity limits viral replication within the tumor, it enhances the antitumor efficacy of polio virotherapy. It shows that the antitumor effect of vaccine-specific recall is contingent upon CD4^+^ T cells; does not require direct infection of, or viral antigen presentation by, malignant cells; and mediates antitumor efficacy through both type I and II immunity. Thus, pre-existing polio immunity is an asset to intratumor polio virotherapy that contributes a lysis-independent mechanism of action.

Given the eminent importance of CD4^+^ T cells in cancer immunotherapy^31,36^, growing efforts aim to leverage CD4^+^ T cell help within tumors, e.g. with CD40 agonistic antibodies^41^, or with peptide vaccines including MHC class II epitopes^10^ for next generation cancer vaccine designs that prime neoantigen specific CD4^+^ and CD8^+^ T cells^51^. Our work uncovers the potential of harnessing childhood vaccine-specific memory CD4^+^ T cells to engage multifaceted antitumor mechansims of CD4^+^ T cells.

## Supporting information

Merged Supplement

## Acknowledgements

We thank G. Palmer (Duke University, NC) for providing the E0771 cell line, M. Mohme (University of Hamburg Medical Center, Hamburg, Germany) for technical advice, and Satoshi Koike (Tokyo Metropolitan Institute of Medical Science, Japan) for hCD155-tg C57BL/6 mice.

## Notes

Funding: PHS: F32CA224593 (M.C.B.), K99CA263021 (M.C.B.), R00CA263021 (M.C.B.), R01NS108773 (M.G. and S.K.N.), R35CA225622 (D.D.B.); Department of Defense Breast Cancer Research Program award W81XWH-16-1-0354 (S.K.N.), National Cancer Center Breast Cancer Project Grant (M.C.B.).

### Competing Interest Statement

M.C.B., D.M.A., D.D.B., S.K.N., and M.G. own intellectual property related to PVSRIPO, which has been licensed to Istari Oncology, Inc. M.C.B, M.G., and D.D.B received consultancy fees from Istari Oncology, Inc.; M.G. and D.D.B hold equity in Istari Oncology, Inc. S.K.N., M.C.B., D.D.B., and M.G. are inventors on patent application PCT/US2017/039953 held/submitted by Duke University that covers the composition and methods for activating antigen presenting cells with PVSRIPO. All other authors declare they have no competing interests.

### Summary of Updates

Figures 1 and 2 were revised for clarity, the main text was updated.

