## Supplementary material for "Intratumor Childhood Vaccine-Specific CD4^+^ T cell Recall Coordinates Antitumor CD8^+^ T cells and Eosinophils": Merged Supplement

Brown, MC et al., 2022

**This file contains:**

**Supplementary Figures 1-11**

**Supplementary methods**

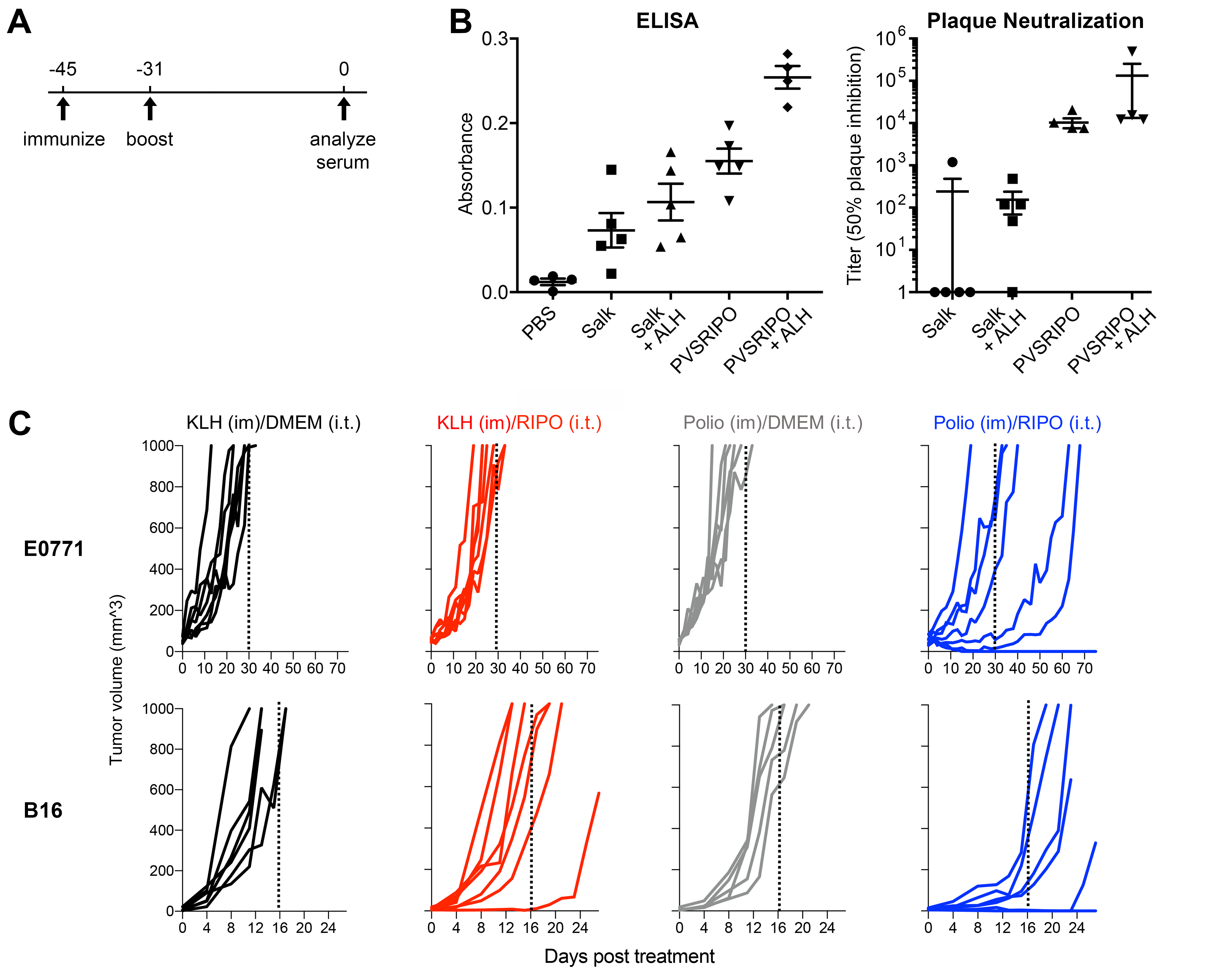

**Supplementary Figure 1.** (**A**) Schema for immunization validation in (B). (**B**) ELISA (left) and plaque neutralization assay (right) from serum of mice immunized as shown (n= 5/group). (**C**) Individual tumor volumes from experiments shown in Figure 1B.

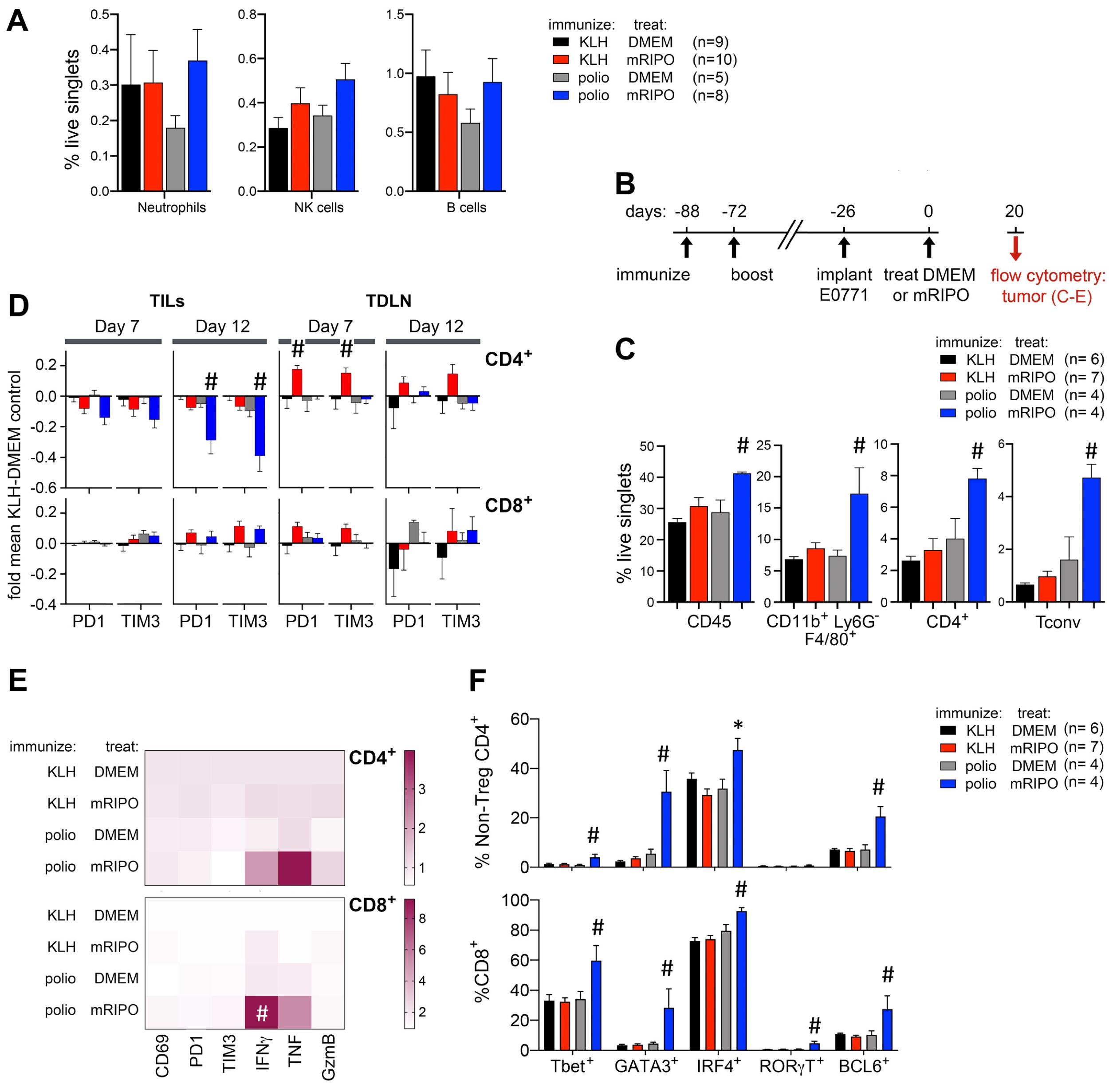

**Supplementary Figure 2.** (**A**) Density of neutrophils, NK cells, and B cells from the experiment presented in Figure 1E. (**B-C**) Experimental schema and analyses of infiltrating immune cells in the E0771 model, as done in Figures 1G-H for B16. (**D**) PD1 and TIM3 surface staining on CD4+ and CD8+ T cells from tumor (TILs) or tumor draining lymph node (TDLN) derived single cell suspensions using the same samples analyzed in Figure 2 at day 12 post treatment, values were normalized to log(fold mean KLH-DMEM control). (**E, F**) Analysis of TIL phenotypes in the E0771 model for experiment presented in (B-C), as done for the B16 model in Fig 2B. (**A, C, D, F**) Data bars represent mean + SEM; (**E**) values were normalized as fold mean KLH-DMEM for each marker; (**C-F**) asterisks denote Dunnett’s multiple comparison test vs corresponding DMEM treated controls (p<0.05, two tailed); # indicates significant Tukey’s post-hoc test vs all other groups.

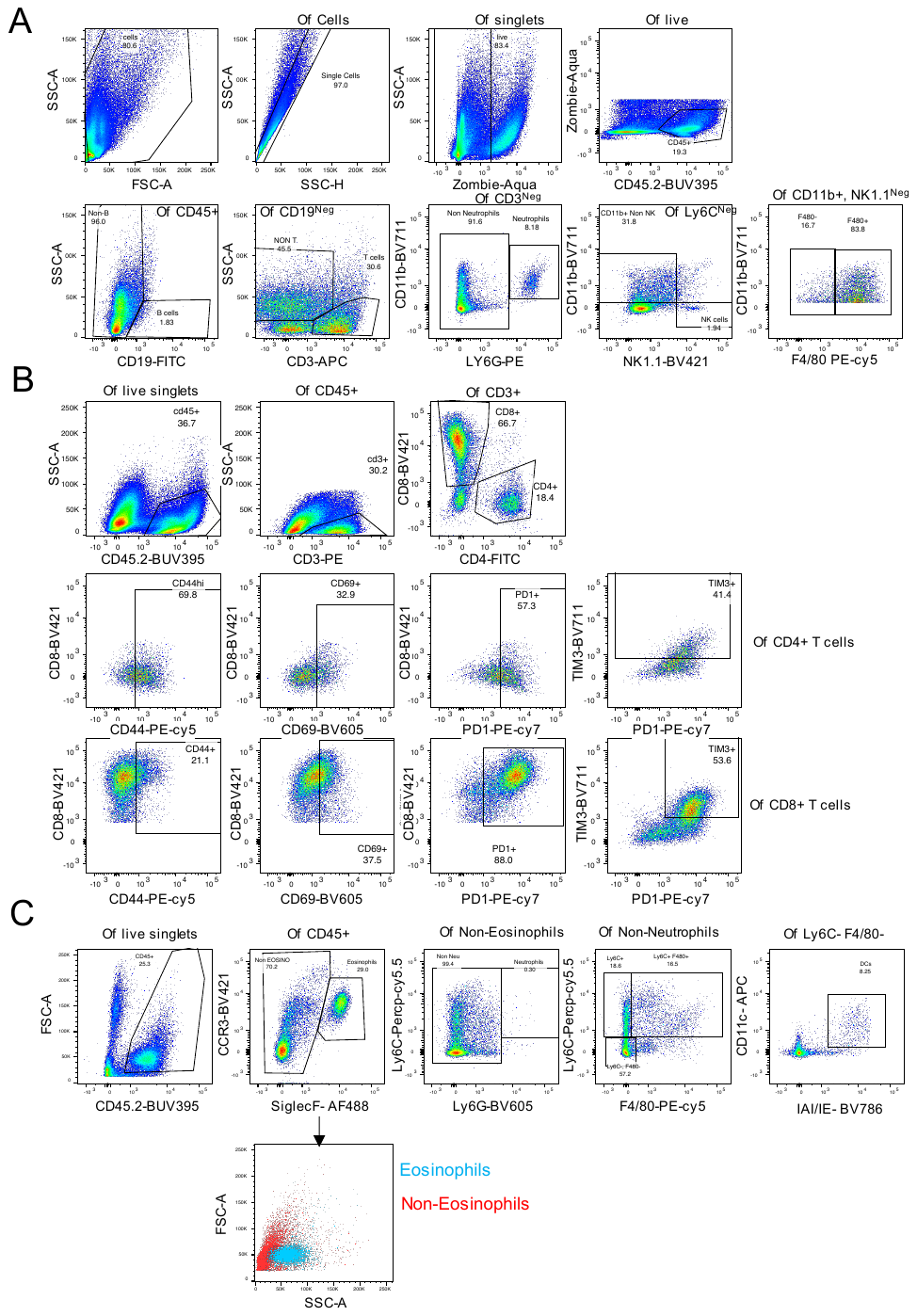

**Supplementary Figure 3.** Representative gating strategy for tumor infiltrating cells analyzed in Figures 1 and S2.

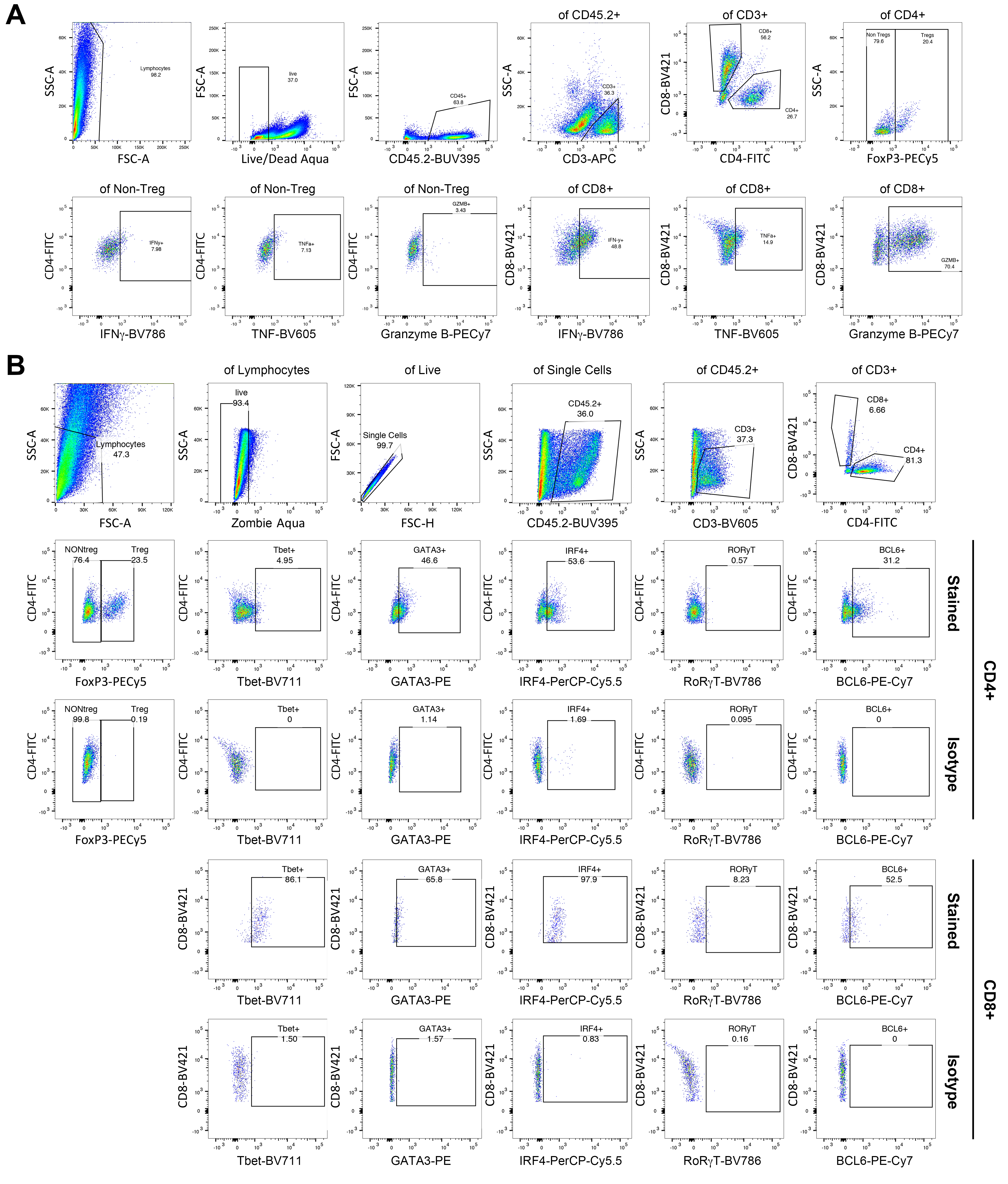

**Supplementary Figure 4.** Representative gating strategy for TIL activation/differentiation status as shown in Figures 2B and S2.

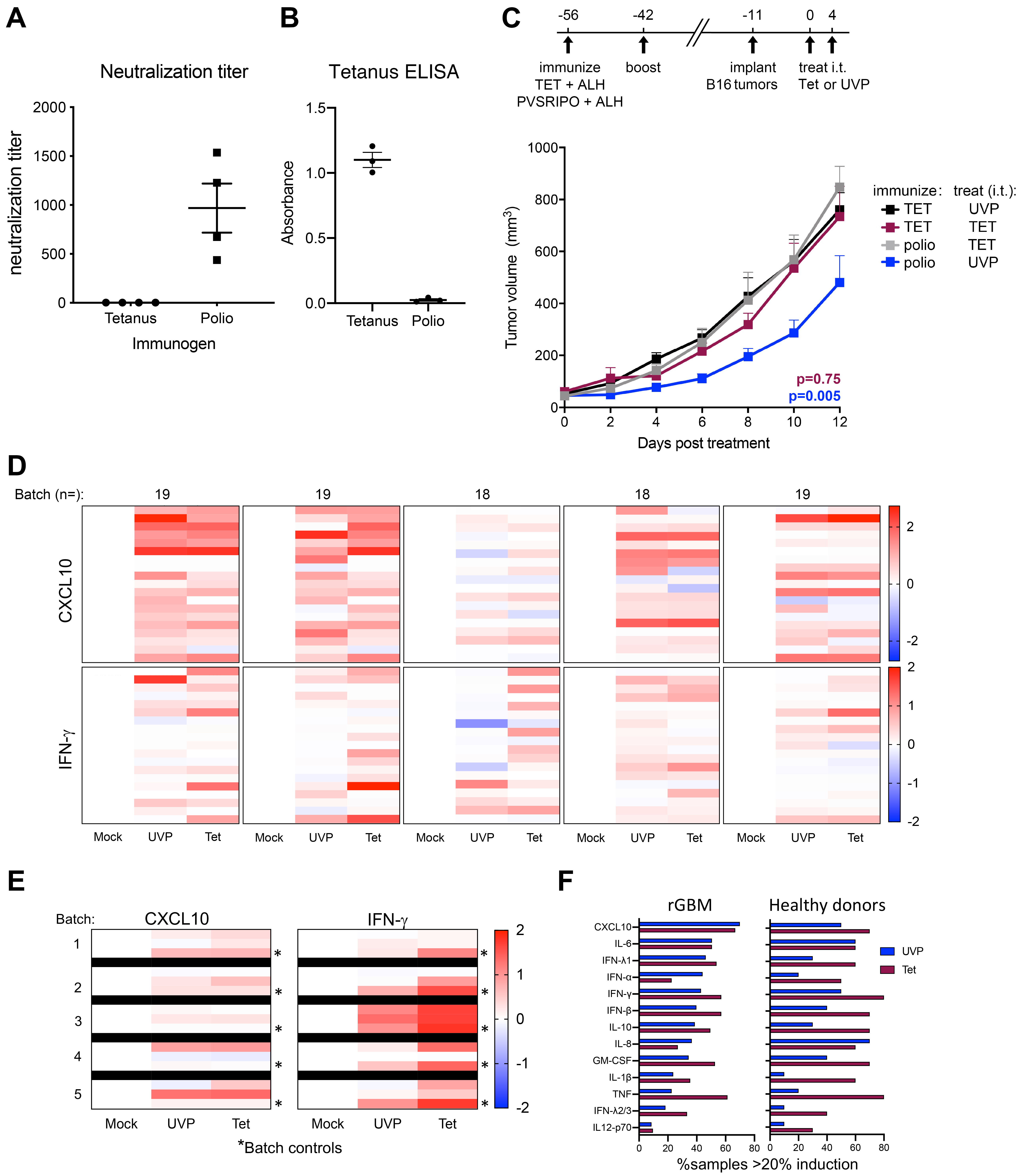

**Supplementary Figure 5.** (**A, B**) Validation of immunogen-specific serum antibodies for immunization strategies in Figure 2A, B, and D by PVSRIPO plaque neutralization assay (**A**; n= 4/group) and tetanus ELISA (**B**; n= 3/group); mean -/+ SEM is shown. (**C**) Repeat tumor therapy experiment for Figure 2A. Mean tumor volume + SEM are shown (Polio-Tet and Tet-UVP: n= 7; Tet-Tet: n= 6; Polio-UVP: n= 8). (**D**) Individual patient-level PBMC CXCL10 and IFN-γ responses for recurrent GBM patients shown in Fig 2E after UVP or Tet treatment; heatmaps are separated by experimental batch and represent log(fold Mock control). (**E**) Individual healthy donor and batch control (*) for CXCL10 and IFN-γ responses to UVP and Tet treatment, normalized as in (D). (**F**) Comparison of the percentage of donors with >20% induction of each presented cytokine relative to Mock treatment for rGBM (left, n=93) and healthy donor (right, n=10) patients after UVP or Tet treatment.

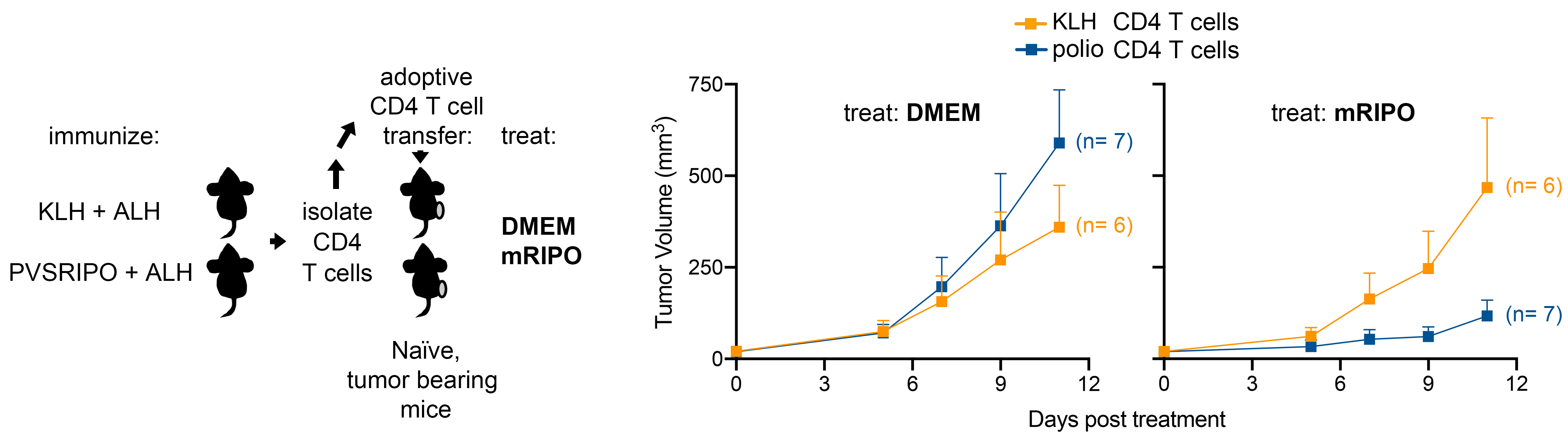

**Supplementary Figure 6.** Repeat experiment of the assay described in Figure 3E, using CD4^+^ T cells from KLH or polio immunized mice. Mean tumor volume + SEM are shown.

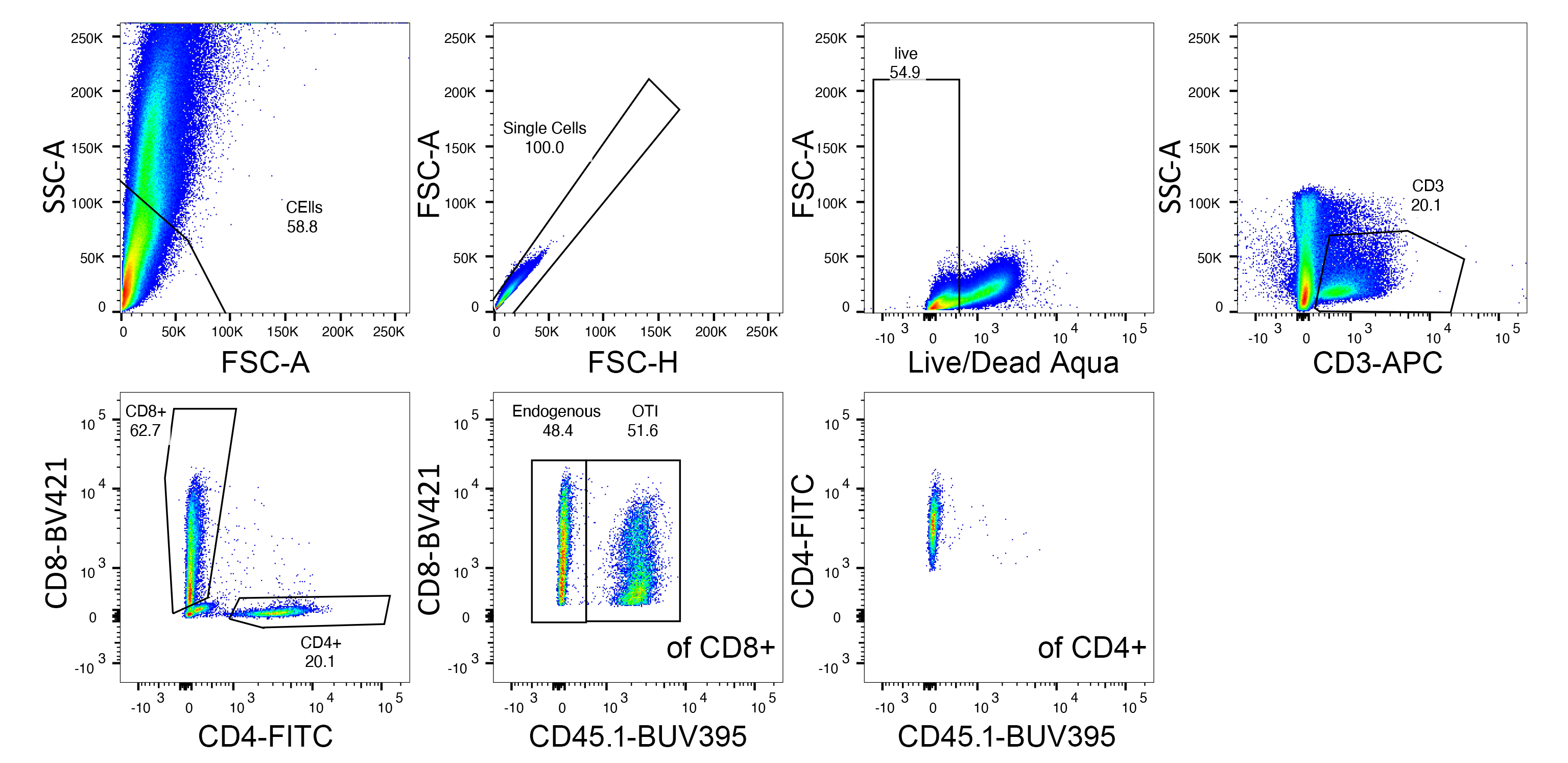

**Supplementary Figure 7.** Gating strategy for OT-I (CD45.1^+^) vs endogenous (CD45.1^Neg^) CD8^+^ T cells in tumors from experiments presented in Figures 4 and 6; gating for markers of activation and differentiation were conducted as shown in Figure S4.

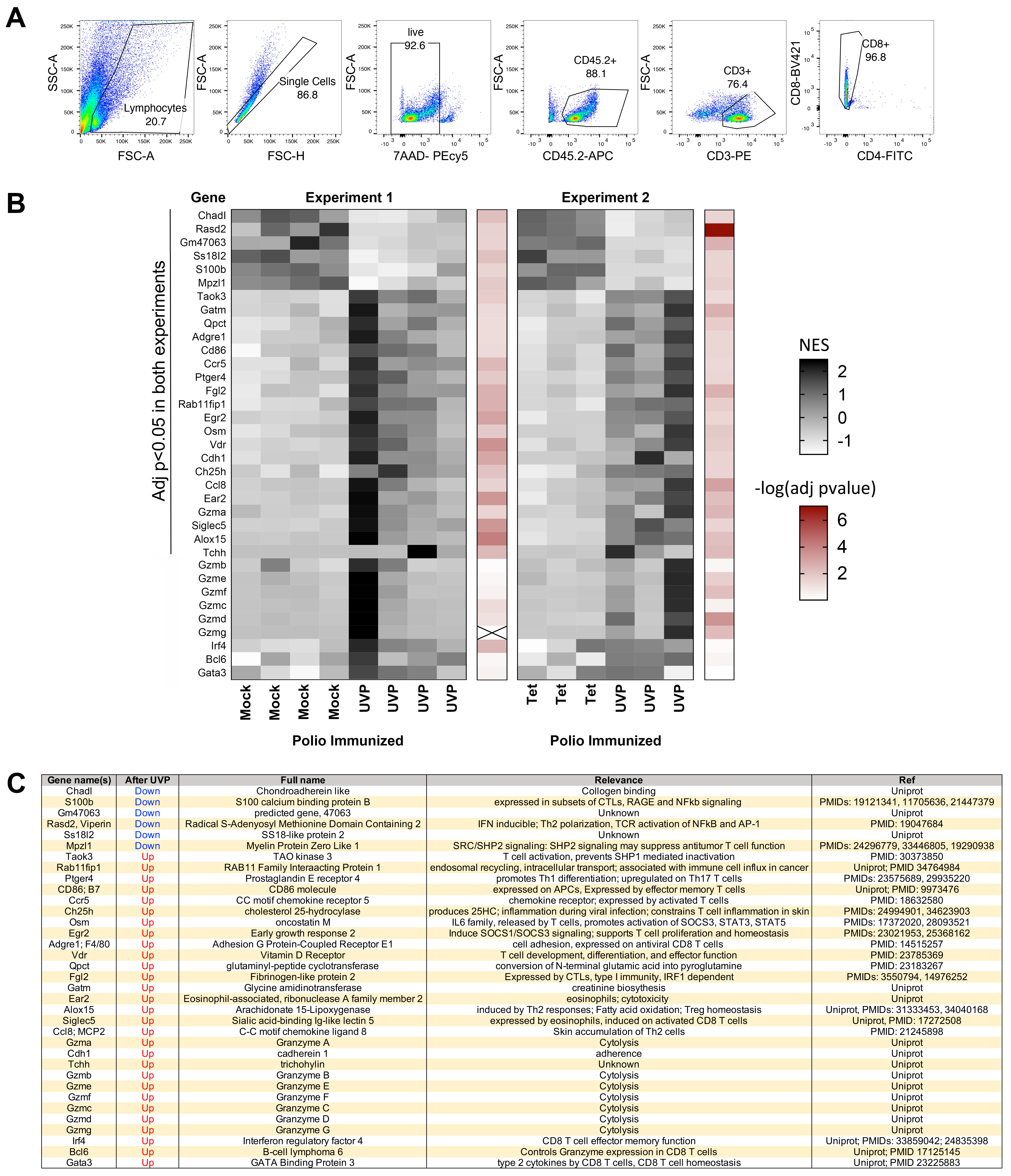

**Supplementary Figure 8.** (**A**) Validation of CD45.1+ OTI+ enrichment after positive CD45.1

selection using tumor suspensions in experiments in Fig 5C; note that OT-I T cells are positive for both CD45.2 and CD45.1, while endogenous T cells were only CD45.2+. (**B**) Normalized enrichment score (NES; black) and -log(p-value) (red) for genes significantly different between OT-I T cells after polio recall (polio immunized mice treated with UVP) vs Mock or Tet treatment in polio immunized mice in two independent experiments as indicated by annotation in figure. In addition, relevant transcripts with protein changes in flow cytometry analyses (Fig 5C): granzymes, IRF4, BCL6, and GATA3 that approached significance in both studies were included; a p-value for Gzmg was not assigned in experiment 1 due to an outlier. (**C**) Table presents known relevance of genes from (B) to T cell biology (PMID= Pub Med ID) and directionality for change (consistent in both replicates) after treatment with UVP in polio immunized mice from two independent experiments.

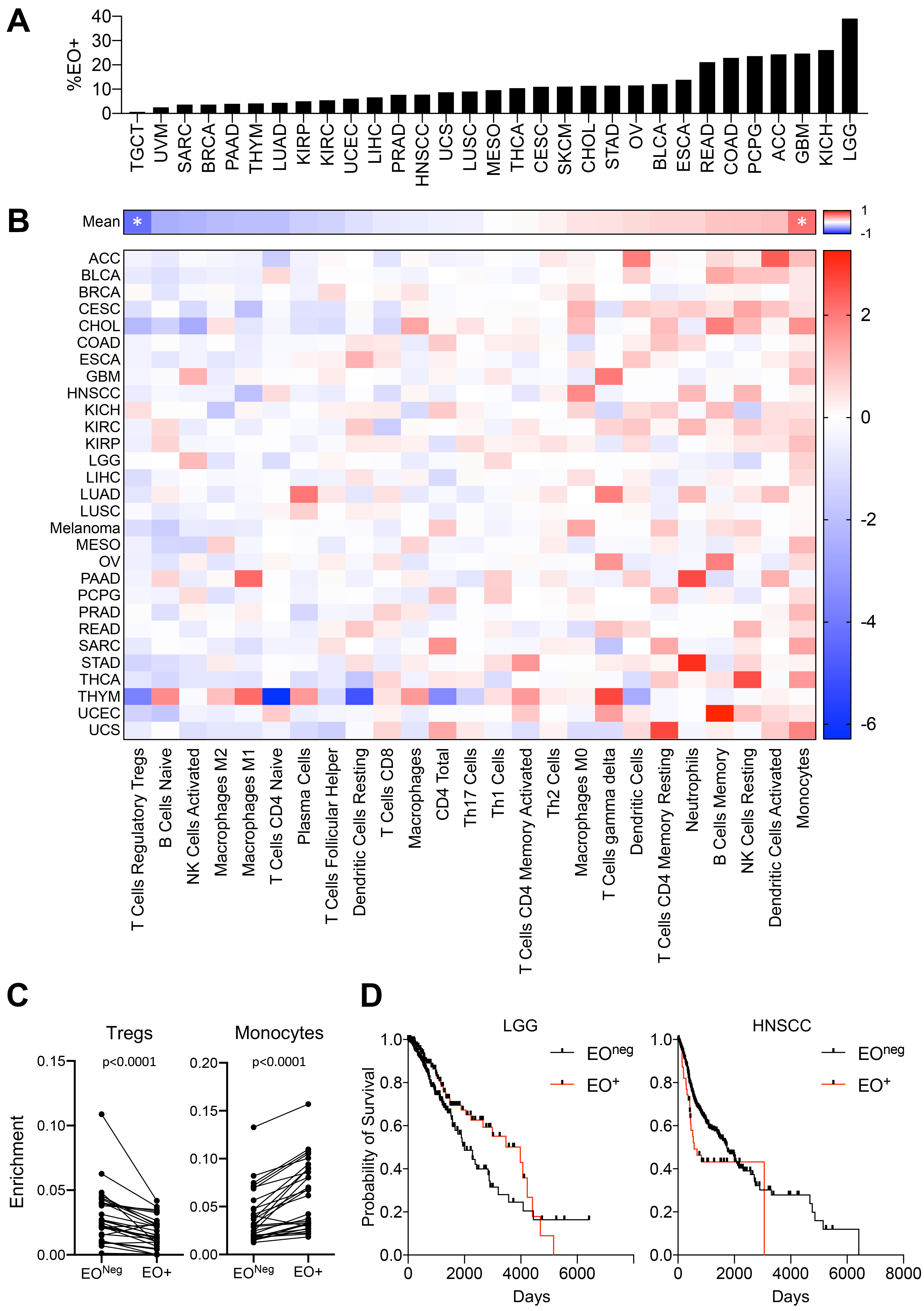

**Supplementary Figure 9.**  Pan-cancer analysis of frequency of eosinophil detection by CIBERSORT within each cancer type (**A**); changes in the frequency of cell types associated with the presence vs absence of detected eosinophils (**B, C**); and survival of LGG and HNSCC by eosinophil detection vs absence corresponding to Figure 6A-B (**D**). (**B**) Asterisks denote significant False Discovery Rate Adjusted paired t-test comparing cell type densities across cancer types (n=29); (**C**) p-values are from paired t-test for monocytes and Tregs across cancer types (n=29); (**B, C**) each cancer type was treated as one pair.

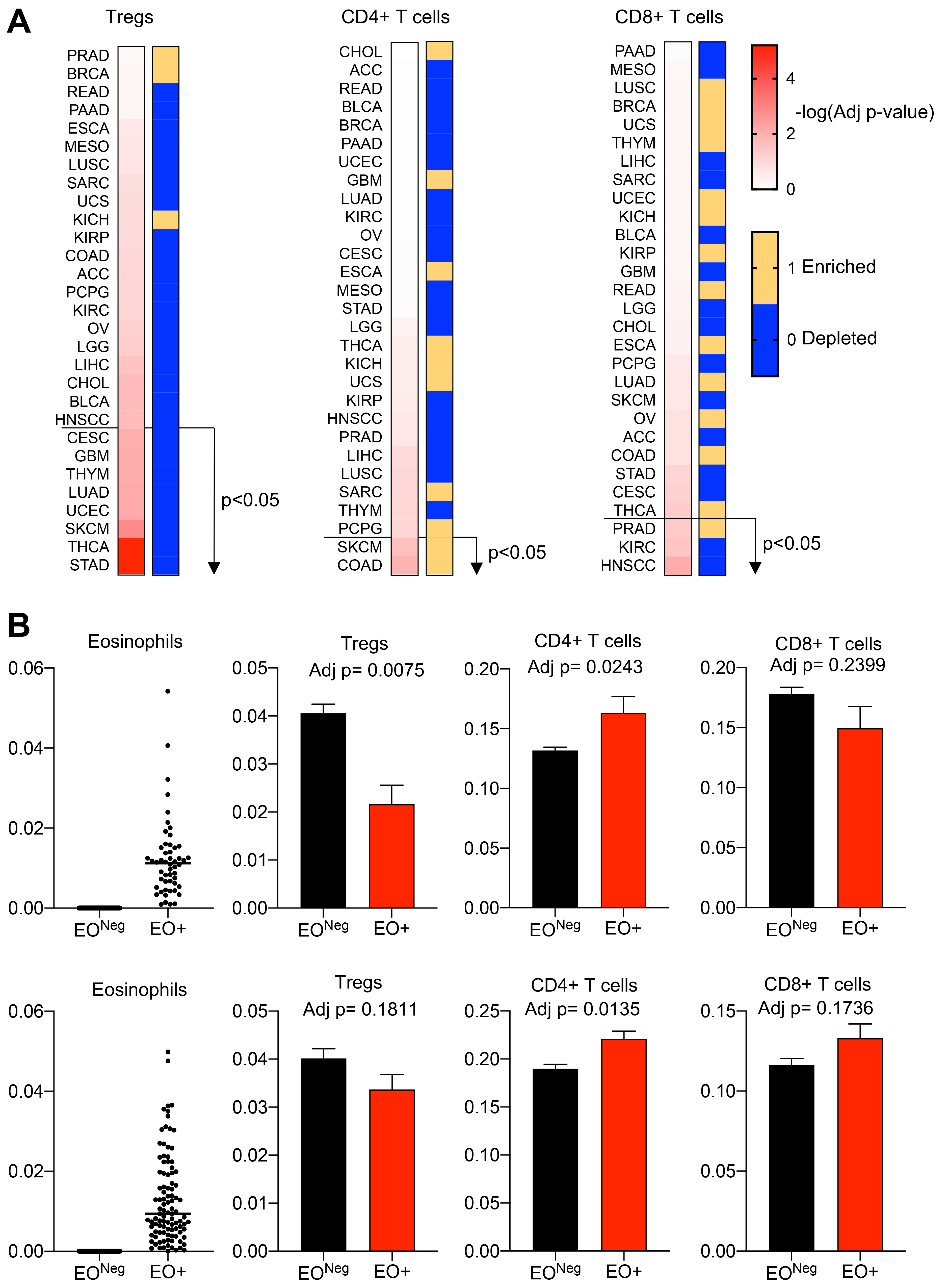

**Supplementary Figure 10.** (**A**) Pan-cancer analysis conducted in Fig 6A and Supplementary Figure 9, presenting False Discovery Rate corrected -log(p-value) and directionality (blue= depleted, yellow= enriched) within each cancer type for Tregs, CD4^+^ T cells, and CD8^+^ T cells; stratified by p-values (high to low) for difference in Tregs. (**B**) Eosinophil, Treg, CD4^+^ T cell, and CD8^+^ T cell predictions for melanoma (SKCM, top) and colorectal cancer (COAD, bottom) for cases with eosinophil detection (EO+) or no eosinophil detection (EO^Neg^). Adj p-values are from False Discovery Rate corrected paired t-test.

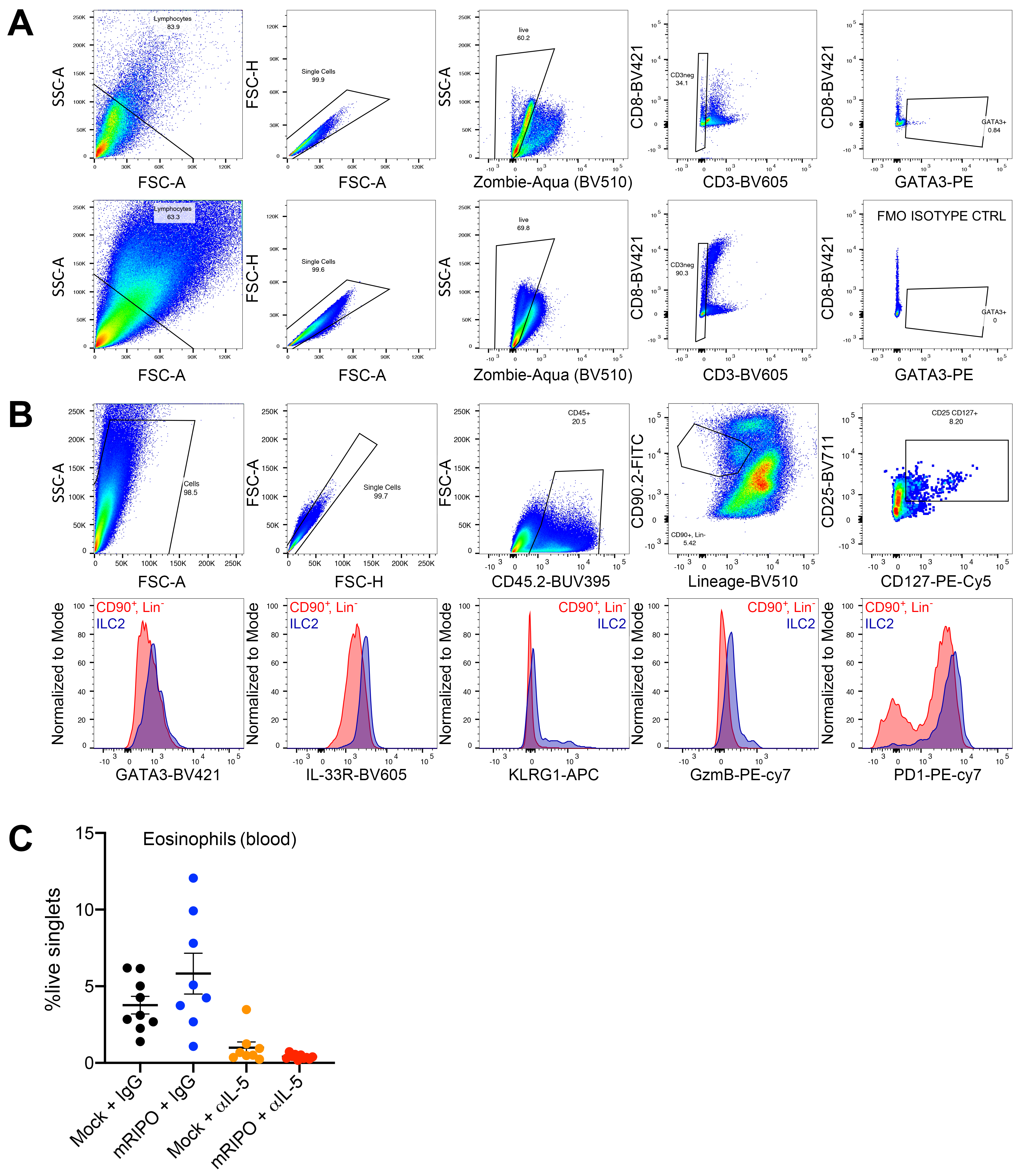

**Supplementary Figure 11.** **(A)** Gating strategy for %GATA3+ CD3^Neg^ cells presented in Figure 5C from prior experiments conducted in Figure 4. FMO= florescence minus one. (**B**) Gating strategy for ILC2s in Figures 5G and 6B, D (top), along with representative histograms showing ILC2 vs Lin^-^CD90.2^+^ cells (bottom). Lineage= Zombie Aqua (live dead), CD3, CD5, CD19, NK1.1, CD11c, CD11b, FCεRIα, γδTCR, αβTCR. (**C**) Blood eosinophil levels determined for the experiment presented in Figure 5D-G.

**SUPPLEMENTARY METHODS:**

*Polio neutralizing antibody assay and ELISA.*

Polio neutralization assays were performed as previously described^51^. ELISA to measure anti-polio antibodies was performed using Maxisorp plates (Nunc) coated with 1x10^7^ pfu PVSRIPO; blocked in PBS + 2% BSA + 0.05% Tween-20 (Sigma-Aldrich); incubated with serially diluted sera (1:20, 1:100, 1:500) for 2h; followed by incubation with 1:30,000 protein-A conjugated HRP (Thermo-Fisher) diluted in PBS for 1h; and development using TMB substrate (Thermo-Fisher). A similar procedure was used for detection of KLH antibodies and Tetanus-specific antibodies, coating plates at a concentration of 10 or 1μg/ml antigen, respectively.

*Flow cytometry antibody panels.*

The panels used in this study include antibodies to the following antigens (purchased from Biolegend unless otherwise noted): lineage panel 1: CD45.2-BUV395 and CD40-BV605 (BD biosciences), Zombie-Aqua Live/dead, NK1.1-BV421, CD11b-BV711, IA/IE-BV786, CD19-FITC, LY6G-PE, F4/80-PE, CD86-PEcy7, CD3-APC; lineage panel 2: CD19-BUV395 (BD biosciences), 7-AAD, CD45.2-APC-Cy7, CD11b-BV711, LY6C-PerCP-Cy5.5, LY6G-PE, CD3-FITC, CD11c-APC; lineage panel 3: CD45.2-BUV395 and CD40-BV605 (BD biosciences), Zombie-Aqua Live/dead, NK1.1-BV421, CD11b-BV711, IA/IE-BV786, CD3/CD19-FITC, LY6G-PE, F4/80-PE, CD86-PEcy7, CD11c-APC; T cell panel: CD45.2-BUV395, Zombie-Aqua Live/dead, CD3-PE, CD4-FITC, CD8-BV421, CD69-BV605, PD1-PE-Cy7 (Thermo-Fisher), TIM3-BV711, CD44-PE-Cy5; Intracellular staining T cell panel 1: CD45.2-BUV395, Zombie-Aqua Live/dead, CD3-APC, CD4-FITC, CD8-BV421, FoxP3-PE-Cy5 (Thermo-Fisher), IFN-γ-BV786, TNF-BV605, Granzyme B-PE-Cy7; intracellular staining T cell panel 2: CD45.2-BUV395, Zombie-Aqua Live/dead, CD3-APC, CD4-FITC, CD8-BV42, FoxP3-PE-Cy5, Tbet-BV711, GATA3-PE, IRF4-PerCP-Cy5.5, RoRγT-BV786, and BCL6-PE-Cy7; OT-I T cell panels were accomplished using the three above T cell panels with exchange of CD45.2 for CD45.1-BUV395; Eosinophil panel: CD45.2-BUV395, CCR3-BV421, Zombie Aqua, CD3/CD19/NK1.1-BV510, CD11c-BV605, CD11b-BV711, IA/IE-BV786, Siglec F-AF488, Ly6C-PERCP-Cy5.5, CD103-PE, F4/80-PECy5, PD1-PEcy7, PD-L1-APC; and ILC2 panel: CD45.2-BUV395, Zombie Aqua, CD3/CD5/CD19/NK1.1/ CD11c/CD11b/FCεRIa/γδTCR/αβTCR-BV510, ST-2(IL33R)-BV605, CD25-BV711, PD1-BV786, CD90.2-AF488, IL5-PE, CD127-PEcy5, Granzyme B-PECy7, KLRG1-APC.

*Associations of eosinophils with other cell types in human tumors and survival.*

CIBERSORT predicted cell type enrichment for each cancer type were obtained from Thorsson et al^36^. Within each cancer type, cases were sorted by eosinophil enrichment, and comparisons were performed between cases with eosinophil score = 0 (EO^Neg^) vs eosinophil score >0 (EO+). Total CD4^+^ T cells reflects the sum of the following scores: ‘T cells CD4 Memory Activated’ + ‘T cells CD4 Memory Resting’ + ‘T cells CD4 Naïve’ + ‘T cells follicular helper.’ For pan-cancer analyses mean enrichment values for each cell type were determined for EO^Neg^ and EO+ for comparison. Mean values from each cell type were scaled and centered within each cell type score across all samples (both EO^Neg^ and EO+), followed by subtracting normalized EO+ from EO^Neg^ values to to generate heatmaps in Fig S9B and Fig S10A. False discovery rate (Benjamini-Hochberg method) was used to adjust for multiple comparisons in Figs S9B and S10A. Hazard ratios and 95% confidence intervals of survival of patients in EO+ vs EO^Neg^ cohorts were determined using the Mantel-Haenszel test and statistical significance was determined using a Mantel-Cox log rank test.
